# Metformin rescues muscle function in BAG3 myofibrillar myopathy models

**DOI:** 10.1101/574806

**Authors:** Avnika A. Ruparelia, Emily A. McKaige, Caitlin Williams, Keith E. Schulze, Margit Fuchs, Viola Oorschot, Emmanuelle Lacene, Meregalli Mirella, Emily C. Baxter, Yvan Torrente, Georg Ramm, Tanya Stojkovic, Josée N. Lavoie, Robert J. Bryson-Richardson

**Affiliations:** School of Biological Sciences, Monash University, Melbourne 3800, Australia; Monash Micro Imaging, Monash University, Melbourne 3800, Australia; Centre de Recherche sur le Cancer de l’Université Laval, Ville de Québec, Quebec, Canada.; Oncologie, Centre de Recherche du Centre Hospitalier Universitaire (CHU) de Québec-Université Laval, Ville de Québec, Quebec, Canada.; Monash Ramaciotti Centre for Structural Cryo-Electron Microscopy, Monash University, Melbourne 3800, Australia; APHP, Centre de Référence de Pathologie Neuromusculaire Nord/Est/Ile-de-France, Institut de Myologie, Laboratoire de Pathologie Risler, GH Pitié-Salpêtrière, Paris, France.; Stem Cell Laboratory, Department of Pathophysiology and Transplantation, Università degli Studi di Milano, Fondazione IRCCS Ca’ Granda Ospedale Maggiore Policlinico di Milano, Centro Dino Ferrari, via F Sforza, 35, 20122 Milan, Italy; Biochemistry and Molecular Biology, Monash Biomedicine Discovery Institute, Monash University, Melbourne, Victoria, Australia.; Centre de référence des maladies neuromusculaires, Hôpital Pitié-Salpétrière, Assistance-Publique Hôpitaux de Paris, Paris, France.; Département de Biologie Moléculaire, Biochimie Médicale et Pathologie, Université Laval, Ville de Québec, Quebec, Canada.

## Abstract

Dominant *de novo* mutations in the co-chaperone BAG3 cause a severe form of myofibrillar myopathy, exhibiting progressive muscle weakness, muscle structural failure, and protein aggregation.

To identify therapies we generated two zebrafish models, one conditionally expressing BAG3^P209L^ and one with a nonsense mutation in *bag3*. Whilst transgenic BAG3^P209L^ expressing fish display protein aggregation, modelling the early phase of the disease, *bag3*^−/−^ fish demonstrate impaired autophagic activity, exercise dependent fibre disintegration, and reduced swimming activity, consistent with later stages.

We confirmed the presence of impaired autophagy in patient samples and screened autophagy promoting compounds for their effectiveness at removing protein aggregates, identifying nine including Metformin. Further evaluation demonstrated Metformin is not only able to remove the protein aggregates in zebrafish and human myoblasts but is also able to rescue the fibre disintegration and swimming deficit observed in the *bag3*^−/−^ fish. Therefore, repurposing Metformin provides a promising therapy for BAG3 myopathy.

## Introduction

Myofibrillar myopathies are a group of chronic muscle diseases characterised at the cellular level by accumulation of protein aggregates and structural failure of the muscle fibre. There is significant variability in the presentation of these diseases, with onset ranging from infantile to late seventies and muscle weakness ranging from mild reductions to severe impairment of skeletal, cardiac, and respiratory muscles resulting in early death. Causative mutations for myofibrillar myopathies have been identified in 10 genes; desmin ^1^, αB-crystallin ^2^, myotilin ^3^, Z-band alternatively spliced PDZ motif-containing protein ^4^, filamin C (FLNC) ^5^, bcl-2 related athanogene 3 (BAG3) ^6^, four and a half LIM domain 1 ^7^, titin ^8^, DNAJB6^9^, and small heat shock protein 8 (HSPB8) ^10^. All of these genes encode proteins found at the Z-disk, a key structure involved in the transmission of tension and contractile forces along the muscle fibre.

While structural failure of the muscle fibre is a feature of myofibrillar myopathies, not all of the proteins associated with the disease have a direct structural role. One such protein is BAG3, a multi domain co-chaperone that is predominantly expressed in skeletal and cardiac muscle, where it co-localises with FLNC and α-actinin at the Z-disk ^11^. In muscle, BAG3 regulates several cellular processes including inhibiting apoptosis and promoting cell survival ^12,13^, regulating protein turnover by stimulating autophagy and inhibiting proteosomal degradation ^14–19^, and regulating cellular mechanotransduction ^18^. Given such important functions of BAG3 it is unsurprising that mutations in it result in disease with two mutations identified to cause myofibrillar myopathy ^6,20^ and nine in dilated cardiomyopathy ^21–24^

The dominant *de novo* myofibrillar myopathy causing BAG3^P209L^ mutation is not only the first identified but also the most well characterized BAG3 mutation ^6,20^. This is the most severe form of myofibrillar myopathy described, patients presenting with early onset (6-8 years of age) and rapidly progressing limb and axial muscle weakness that is often followed by cardiomyopathy, respiratory failure, and/or neuropathy ^6,20^. At a cellular level abnormal accumulation of sarcomeric and extracellular matrix proteins is evident, along with presence of granular-filamentous aggregates, Z-disk thickening, and myofibrillar disintegration ^6,20,25^.

To examine the mechanistic basis of BAG3 myofibrillar myopathy we previously generated zebrafish models that transiently express BAG3^P209L^ in a subset of muscle cells, or have reduced *bag3* expression ^26^. Using these models, we demonstrated that whilst the protein aggregation is due to the presence of BAG3^P209L^, the structural failure of the muscle fibre is due to protein insufficiency following sequestration of both wildtype and mutant protein in the aggregates. Our data suggested that strategies that promote the clearance of aggregates, and subsequently prevent the depletion of functional BAG3 may be highly beneficial for the treatment of myofibrillar myopathy ^26^. The identification of treatments for rare disease, such a BAG3 myopathy, are a challenge as the small patient population limits the studies that may be conducted on novel compounds. As such, in the current study we explored the potential of known autophagy inducing compounds, many of which are already approved by the US Food and Drug Administration (FDA) for clinical use, as potential therapies for BAG3 myofibrillar myopathy.

We established zebrafish models for BAG3 myofibrillar myopathy and completed a screen of known autophagy promoting drugs identifying several effective at removing protein aggregates. We further identified a defect in autophagy in these fish, which was subsequently confirmed in patients. Our study suggests treatment of BAG3 myofibrillar myopathy would be achieved through stimulation of autophagy, removing protein aggregates and overcoming defects in autophagy.

## Results

### Expression of BAG3^P209L^ results in the formation of protein aggregates

To determine the consequence of BAG3^P209L^ expression we generated zebrafish conditionally expressing fluorescently tagged, full-length, human wildtype BAG3 (Tg (BAG3^wt^-eGFP)) or BAG3^P209L^ (Tg (BAG3^P209L^-eGFP)) in skeletal muscle (Figure 1A). Both transgenes are expressed at comparable levels in the zebrafish myotome (Figure 1B). At 48 hours post fertilisation (hpf) whilst both BAG3^wt^-eGFP and BAG3^P209L^-eGFP localize to the Z-disk of the sarcomere (Figure 1B, Supplementary Figure 1), BAG3^P209L^-eGFP additionally localizes to the myosepta and forms protein aggregates throughout the muscle cell. By 144 hpf the BAG3^P209L^-eGFP aggregates become more evident at the myoseptal boundaries. Tg (BAG3^P209L^-eGFP) fish therefore recapitulate one of the key hallmark features of the disease, the formation of protein aggregates.

**Figure 1:**
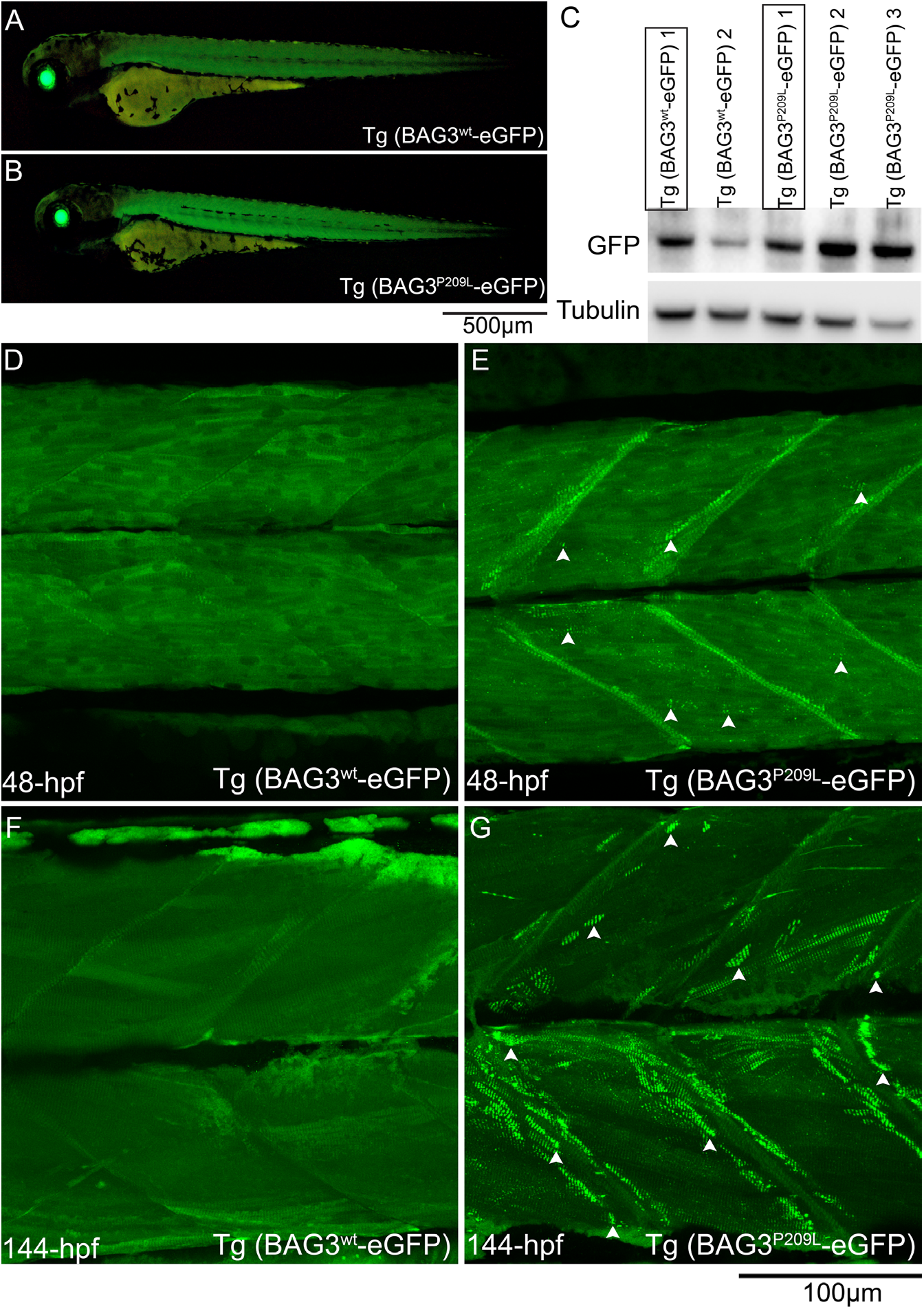
Expression of BAG3^P209L^ results in protein aggregation. (A-B) Live images of 48 hpf transgenic zebrafish expressing BAG3^wt^-eGFP or BAG3^P209L^-eGFP specifically in skeletal muscle, with lens GFP expression indicating presence of Cre protein. (C) Western blot showing differential levels of transgene expression in the multiple Tg(BAG3^wt^-eGFP) and Tg(BAG3^P209L^-eGFP) strains. Strains with comparable transgene expression (boxed) are used in all subsequent experiments. (D-G) Live confocal images of Tg(BAG3^wt^-eGFP) and Tg(BAG3^P209L^-eGFP) embryos at 48-hpf and 144-hpf. Both, BAG3^wt^-eGFP and BAG3^P209L^-eGFP localize to the sarcomere but BAG3^P209L^-eGFP additionally localizes to the myoseptal boundary, and forms protein aggregates (arrow heads) along the length of the muscle fibre, and around the myoseptal boundaries.

### Reduced Bag3 results in contraction dependent fibre disintegration

Given the hypothesis that protein aggregates ultimately lead to BAG3 loss of function, using CRISPR/Cas9 genome editing we generated a *bag3* mutant with a 41 base pair insertion in exon 2, resulting in a frameshift and incorporation of a premature stop codon, and subsequent nonsense mediated decay of the mRNA (Figure 2A). Whilst under normal conditions the muscle structure in *bag3* heterozygous (*bag3*^+/−^) and homozygous mutants (*bag3*^−/−^) is indistinguishable from that of wildtype embryos (*bag3*^+/+^), *bag3*^+/−^ and *bag3*^−/−^ embryos display a structural failure of the myofibre following incubation in methyl cellulose, a viscous solution that increases the load on the muscle Figure 2B, 2C, and 2D). A reduction in, or loss of Bag3, therefore results in contraction dependent fibre disintegration, which is the second defining feature of myofibrillar myopathy.

**Figure 2:**
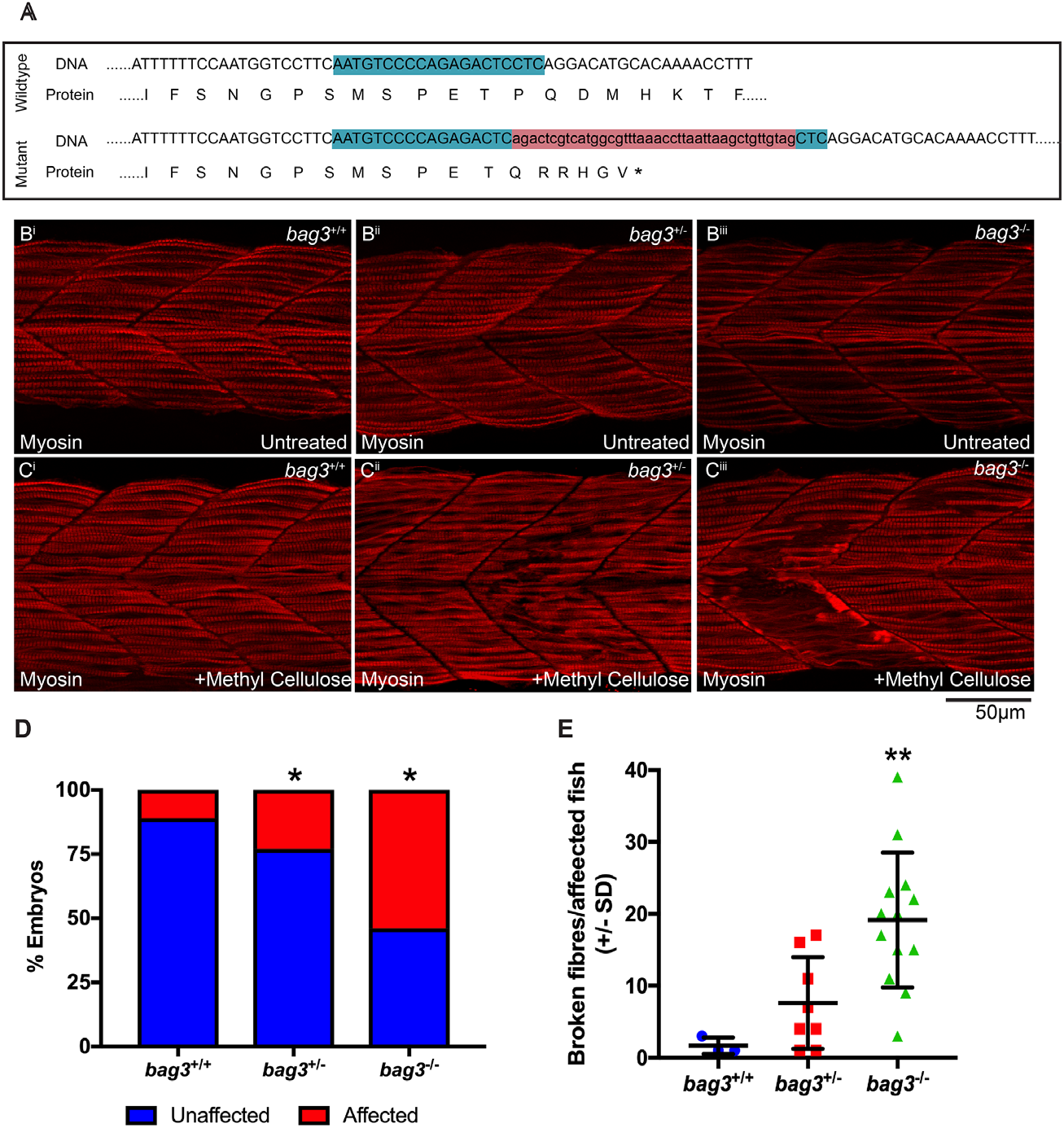
Bag3 deficient zebrafish embryos display fibre disintegration. (A) Schematic of Bag3 wildtype and mutant exon 2 genomic DNA sequence, and resulting protein, with the mutant predicted to incorporate a premature stop (*). The mutant was generated using CRIPSR/Cas9 genome editing. The CRISPR target site is highlighted in green and the 41bp insert in red. (B) Muscle fibres span the entire length of the somite in the 26 hpf wildtype (*bag3*^+/+^), Bag3 heterozygous (*bag3*^+/−^) and Bag3 mutant (*bag3*^−/−^) embryos as seen by Myosin antibody labelling. (C) Incubation of 26 hpf *bag3*^+/−^ and *bag3*^−/−^ in methyl cellulose results in fibre disintegration, which is not evident in *bag3*^+/+^ embryos. (D) Graph showing the percentage of affected *bag3*^+/+^, *bag3*^+/−^ and *bag3*^−/−^ embryos the latter two genotypes having a significant increase in the proportion of phenotypic fish (p<0.05 as determined using a chi-square test (chi-square = 28.3, df=2). Quantification of the number of broken fibres in affected embryos following incubation in methyl cellulose with *bag3*^−/−^ showing a significant increase – as calculated using a one-way ANOVA (F = 8.697, df = 2, p = 0.0049). Error bars represent +/− SD from three biological replicates. The total number of fish examined in each replicate is documented in Supplementary Table 2. *p<0.05, **p<0.01.

### Reduced Bag3, but not the presence of BAG3^P209L^, results in muscle weakness

To assess the pathogenicity of BAG3^P209L^ and the loss of *bag3* on skeletal muscle, we examined muscle function in both the transgenic overexpression models and the loss of function *bag3*^−/−^ mutant. At 2 dpf we performed touch evoked response assays to measure the maximum acceleration during a burst swimming motion, acceleration being directly proportional to the force produced by the muscle. We reveal no significant differences in the maximum acceleration in Tg (BAG3^wt^-eGFP) (Figure 3A) and Tg (BAG3^P209L^-eGFP) (Figure 3B) when compared to their respective sibling controls, and in *bag3*^−/−^ mutants when compared to *bag3*^−/−^ and *bag3*^+/−^ embryos (Figure 3E).

**Figure 3:**
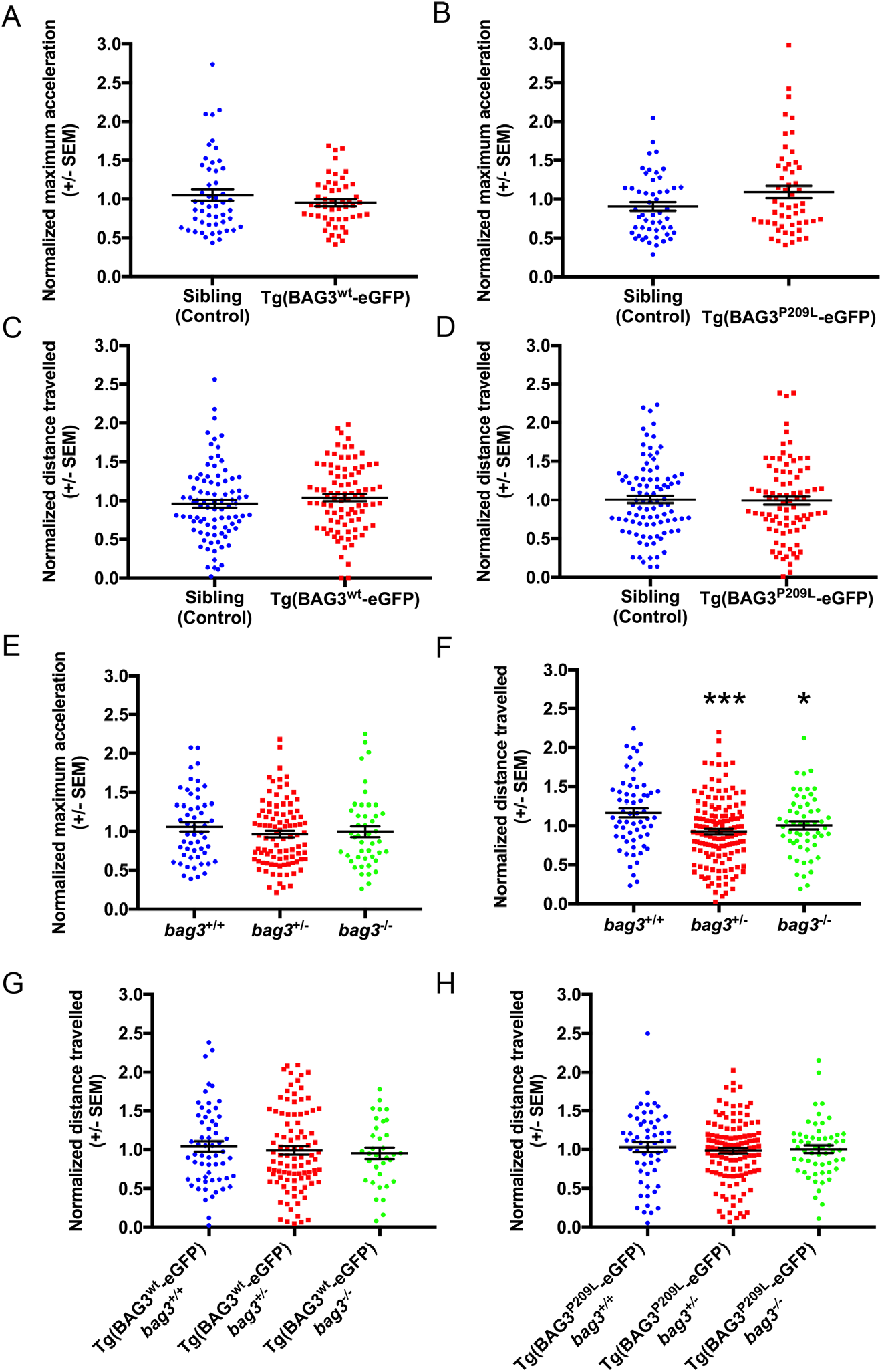
Quantification of muscle function in BAG3 myofibrillar myopathy models. (A-B) Quantification of the maximum acceleration recorded from touch-evoked response assays in 48-hpf Tg(BAG3^wt^-eGFP) and Tg(BAG3^P209L^-eGFP) normalized to their respective sibling controls. (C-D) Quantification of distance travelled by 6-dpf Tg(BAG3^wt^-eGFP) and Tg(BAG3^P209L^-eGFP) normalized to their respective sibling controls. (E) Normalized maximum acceleration of *bag3*^+/+^, *bag3*^+/−^ and *bag3*^−/−^. (F) Normalized distance travelled of *bag3*^+/+^, *bag3*^+/−^ and *bag3*^−/−^. (G-H) Normalized distance travelled by Tg(BAG3^wt^-eGFP) and Tg(BAG3^P209L^-eGFP) on *bag3*^+/+^, *bag3*^+/−^ and *bag3*^−/−^ background. Error bars represent +/− SEM for three-four biological replicates. The number of fish in each replicate is documented in Supplementary Table 3. *p<0.05, ***p<0.01 calculated using a one way ANOVA with Dunnett’s post-hoc multiple comparison correction test.

We also measured, at 6 dpf, the distance swum, which is an indicator of muscle performance. Whilst the distance swum by Tg (BAG3^wt^-eGFP) (Figure 3C) and Tg (BAG3^P209L^-eGFP) (Figure 3D) was comparable to their respective sibling controls, both *bag3*^+/−^ and *bag3*^−/−^ were found to have significant reductions in distance swum (Figure 3F). A 50% reduction or complete loss of *bag3* is therefore sufficient to cause impaired muscle performance.

We next examined the capacity of BAG3^wt^ and BAG3^P209L^ to rescue the muscle impairment observed in loss of function model. Remarkably, expression of BAG3^wt^-eGFP or BAG3^P209L^-eGFP in *bag3*^+/−^ and *bag3*^−/−^ larvae was sufficient to rescue the phenotype resulting from reduction of Bag3 (Figure 3G and 3H). These results not only confirm that the muscle weakness seen in *bag3*^−/−^ larvae is due to loss of Bag3, but they also demonstrate that BAG3^P209L^-eGFP is functional and capable of preventing muscle weakness associated with the loss of Bag3.

### Bag3 mutants have impaired autophagy, potentially explaining their decreased muscle function

Given that fibre disintegration was seen in 24 hpf *bag3*^−/−^ embryos we examined if this structural failure was responsible for the muscle weakness observed at 6 dpf. To visualize the muscle structure, we crossed the Bag3 mutant to a transgenic strain in which GFP tagged FLNC is specifically expressed in muscle. Live imaging of the muscle structure in larvae revealed no gross muscle defects in *bag3*^−/−^ larvae (Supplementary Figure 2A), indicating that fibre disintegration, as seen in the methyl cellulose treated embryos at 24-hpf, is not responsible for muscle weakness in the loss of function model. The 6dpf fish are, however, still susceptible to muscle damage following increased load as 6-dpf, FLNC-eGFP expressing *bag3*^−/−^ larvae incubated in methyl cellulose, display infrequent loss of structural integrity, not observed in *bag3*^+/+^ and *bag3*^+/−^ larvae (Supplementary Figure 2A).

Since we identified no gross defects in the *bag3*^−/−^ larvae not incubated in methyl cellulose, we further examined muscle structure using electron microscopy. Whilst no sarcomeric defects were evident at an ultrastructural level, we observed a four-fold increase in large, electron-dense, double membranous, autophagic vacuoles at the sarcolemma of *bag3*^−/−^ larvae compared to *bag3*^+/+^ larvae (*bag3*^+/+^ : 53 vesicles in 151 muscle cells, average of 0.351 vesicles/ cell; *bag3*^−/−^: 198 vesicles in 134 muscle cells, average of 1.478 vesicles/ cell, Figure 4A and 4B). The structure of the vacuoles seen in *bag3*^−/−^ fish was similar to that of *bag3*^+/+^ fish however, in *bag3*^−/−^ larvae the compartments were more darkly stained (Figure 4C and Figure 4E) and often had protrusions extending into the muscle fibre (Figure 4C). The increased number and increased electron density of autophagic vacuoles in *bag3*^−/−^ larvae suggests an impairment in autophagy following loss of Bag3. We therefore examined autophagic flux in 6 dpf *bag3*^+/+^, *bag3*^+/−^, *bag3*^−/−^ larvae by assessing the difference in LC3 accumulation between untreated and NH_4_Cl treated larvae. We observed a significant reduction in LC3 accumulation in *bag3*^−/−^ compared to *bag3*^+/+^ larvae (Figure 4F and 4G) demonstrating that loss Bag3 results in reduced autophagic capacity, which may contribute to the functional deficit observed following loss of Bag3.

**Figure 4:**
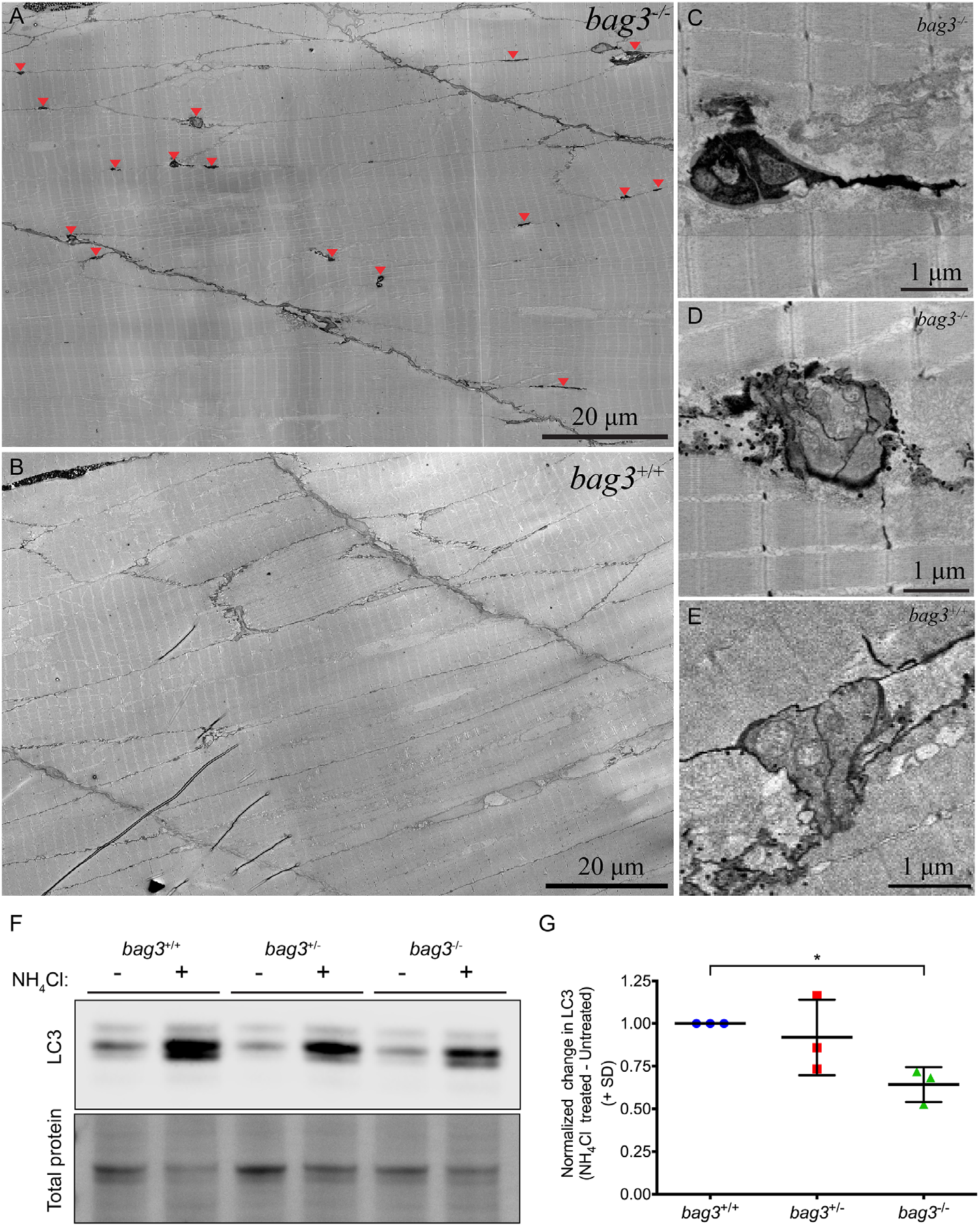
Ultrastructural characterization of muscle in *bag3* deficient zebrafish embryos. Whilst the sarcomeric structure of *bag3*^−/−^ (A) is undistinguishable to that of *bag3*^+/+^ (B), *bag3*^−/−^ additionally have an increased prevalence in sarcolemma associated, double membrane, electron dense autophagic vacuoles (arrow head). (C-D) Autophagic vacuoles in *bag3*^−/−^ embryo are darkly stained, have multiple compartments and occasionally have protrusions extending into the muscle fibre. (E) Autophagic vacuoles in *bag3*^+/+^ embryo have multiple compartments but are not electron dense. (F) Western blot for LC3 and Direct blue stain for total protein (loading control) on protein lysates obtained from *bag3*^+/+^, *bag3*^+/−^ and *bag3*^−/−^ with or without ammonium chloride (NH4Cl) treatment. (G) The autophagic flux determined by normalizing LC3 levels to that of total protein. The change in LC3 levels following NH_4_Cl treatment is presented relative to the levels in the respective untreated groups. Error bars indicate standard deviation. * p<0.05 calculated using a one way ANOVA with Dunnett’s post-hoc multiple comparison correction test (F = 5.288; df =2).

### Myofibrillar myopathy patients also have an impairment in autophagy

Given that we observed impaired autophagy in our zebrafish model of BAG3 myofibrillar myopathy we sought to investigate if autophagy was also impaired in myofibrillar myopathy patients. Western blot experiments revealed accumulation of p62 and LC3 in BAG3 myofibrillar myopathy patient muscle compared to a healthy control (Figure 5A). Additionally, using immunohistochemistry we observed abnormal accumulation of p62 within muscle cells of BAG3^P209L^ patients (Figure 5C), and in myofibrillar myopathy patients carrying mutations in Desmin, Myotilin, or ZASP (Supplementary Figure 3). Taken together, as identified in the zebrafish model, impaired autophagy is a feature of BAG3-related and other myofibrillar myopathies.

**Figure 5:**
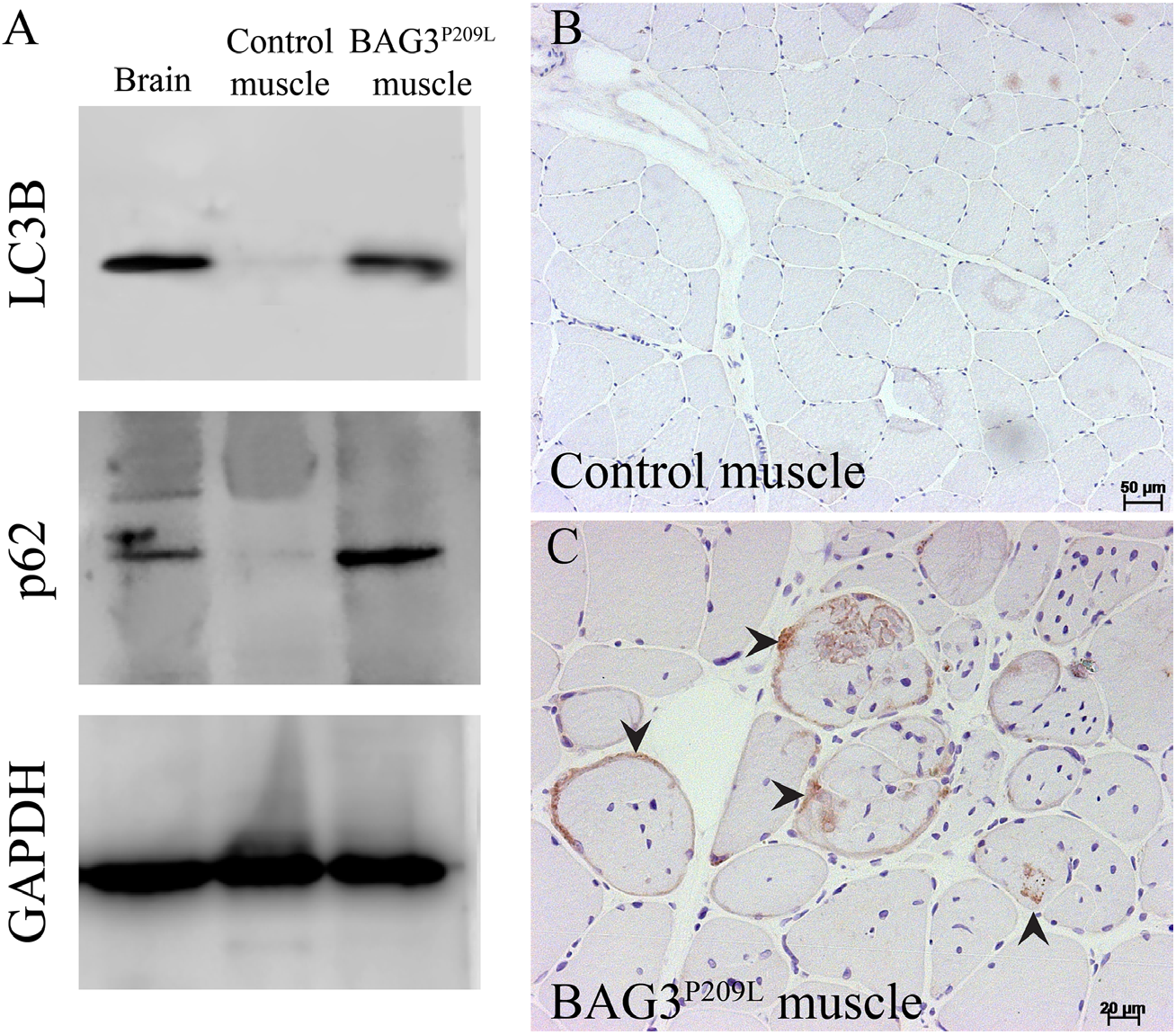
Defective autophagy is evident in BAG3^P209L^ myofibrillar myopathy patients. (A) Western blot for LC3 and p62 showing accumulation in a BAG3^P209L^ patient but not in control muscle. Abnormal p62 accumulation is observed in patient (C) muscle cells but not in control samples (B).

### Increasing autophagy significantly reduces BAG3^P209L^ protein aggregates

Our data suggests that the expression of BAG3^P209L^ and the formation of protein aggregates does not directly cause muscle weakness, with loss of BAG3 due to the sequestration of BAG3 within aggregates resulting in impaired autophagy and subsequent muscle weakness. As such we hypothesized that upregulation of autophagy may be beneficial for the treatment of BAG3 myofibrillar myopathy by not only promoting the clearance of aggregates and preventing any further sequestration of functional protein, but also by compensating for the reduced levels of basal autophagy in affected muscle fibres. We therefore performed a screen of 71 known autophagy promoting drugs to identify the most effective at removing aggregates in our BAG3^P209L^-eGFP expressing fish. 32 hpf Tg (BAG3^P209L^-eGFP) embryos were incubated in one of 71, randomly assigned, drugs or in unsupplemented E3 embryo medium or DMSO supplemented E3 embryo medium. Following 16 hours of drug treatment the embryos were fixed, imaged, and the number of aggregates were quantified using an automated aggregate quantification tool. Eight of the 71 compounds tested were toxic at the dose used and were not further characterized (Supplementary Figure 4). Six compounds, although predicted to increase autophagy, were found to significantly increase the number of aggregates when compared to DMSO control treated embryos (Supplementary Figure 4). We identified nine drugs that were highly effective at reducing aggregate number when compared to DMSO treatment (Figure 6, Supplementary Figure 3).

**Figure 6:**
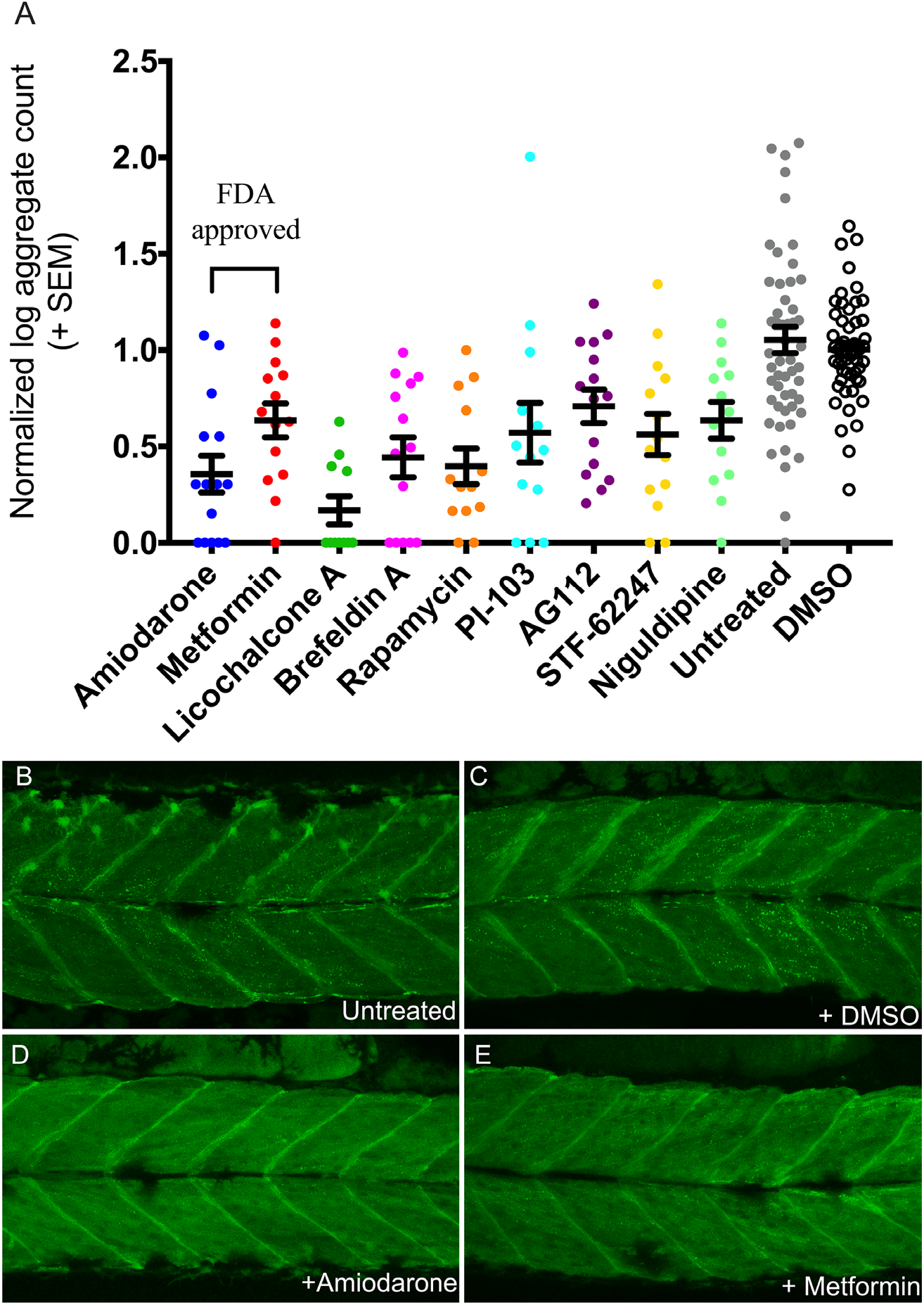
Pharmacological stimulation of autophagy promotes the clearance of aggregates. (A) Graph depicting normalized aggregate number of the nine drugs that significantly (p<0.05) reduce protein aggregation in Tg(BAG3^P209L^-eGFP) embryos, compared to DMSO control. Amiodarone and Metformin are the two drugs currently approved for clinical use by the Federal Drug Association. Representative confocal images of untreated 48-hpf Tg(BAG3^P209L^-eGFP) embryos or treated with DMSO, Amiodarone or Metformin, the latter two resulting in reduced protein aggregation. Error bars represent +/-SEM with three independent biological replicates. The number of fish in each replicate is documented in Supplementary Table 4.

Two of these drugs, Metformin and Amiadarone are currently approved by the FDA, which would allow for rapid translation to clinical use, therefore we further investigated their efficacy in human myoblasts. Similar to the zebrafish experiments, overexpression of BAG3^P209L^-GFP, but not BAG3^wt^-GFP, resulted in protein aggregation (Figure 7A-B). Whilst the majority of BAG3^wt^-GFP localizes to the soluble protein fraction, the majority of BAG3^P209L^-GFP is in the insoluble fraction (Figure 7C). Having shown that BAG3^P209L^ forms protein aggregates in human myoblasts, we examined the ability of Metformin and Amiodarone to reduce them. Whilst Amiodarone treatment had no effect on protein aggregation, Metformin treatment resulted in a siginificant reduction in the number of aggregate containing myoblasts, with the aggregate volume and intensity also significantly reduced (Figure 7D-7I).

**Figure 7:**
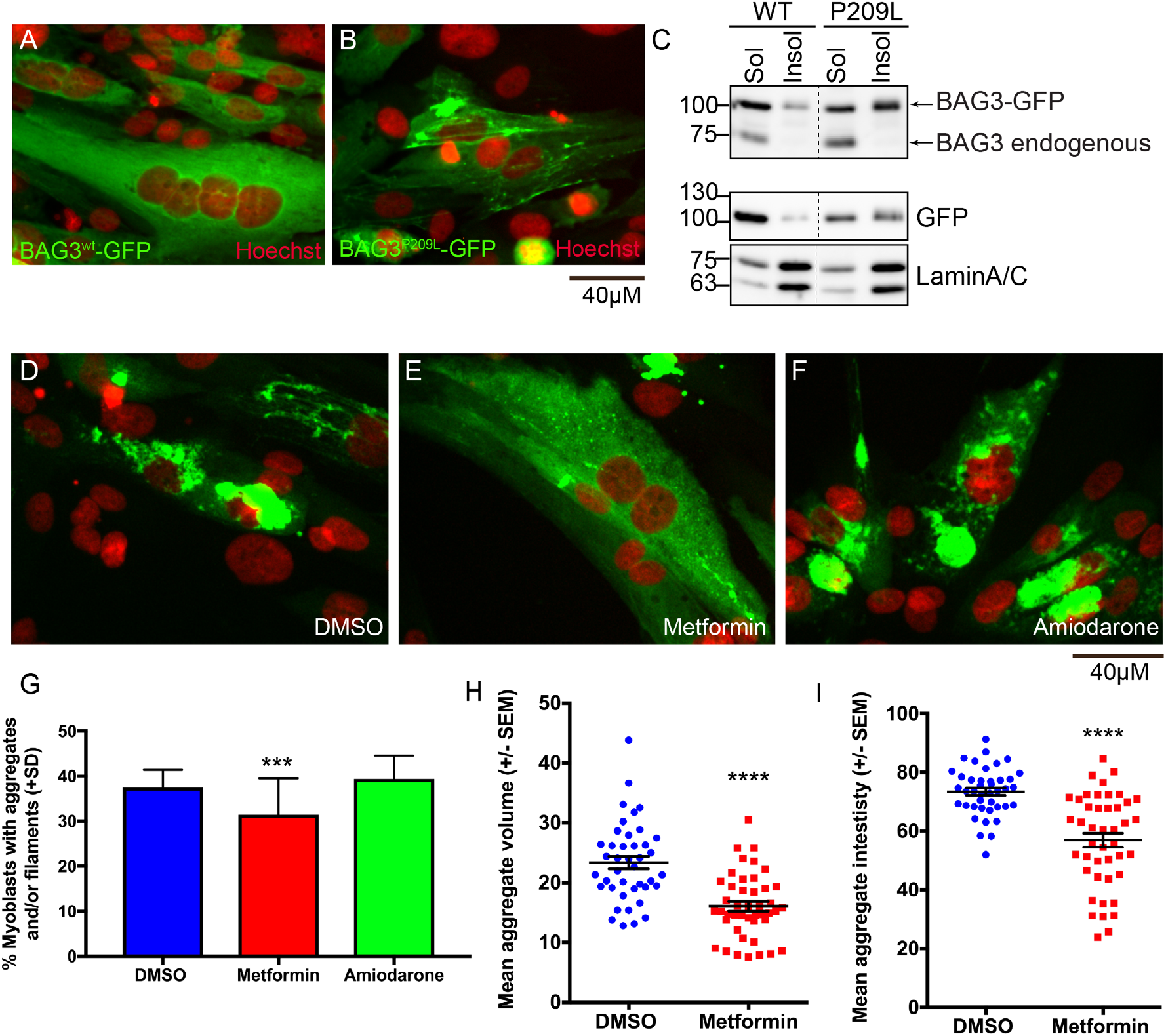
Metformin reduces protein aggregation in human myoblasts. (A-B) Confocal images of human myoblasts transfected with BAG3^wt^-eGFP or BAG3^P209L^-eGFP with protein aggregation evident in the later. (C) Western blot showing that whilst majority of BAG3^wt^-eGFP is found within the soluble lysate, a large proportion of BAG3^P209L^-eGFP is localized to the insoluble fraction. (D-F) Confocal images showing protein aggregation in human myoblasts following 48 hour treatment with DMSO (n=1719), Metformin (n=1664) or Amiodarone (n=970). (G) Metformin but not Amiodarone results in a significant reduction in the percentage of myoblasts with aggregates and/or filaments. ***p<0.001 as determined using a chi-square test (p =0.0001, chi-square = 17.88, df = 2). Metformin (n=44) additionally significantly reduces aggregate volume (H) and intensity (I) compared to DMSO (n=41) treated myoblasts - as determined using a T-test. (****p<0.001, t = 5.561, df = 83).

### Metformin reduces fibre disintegration, and rescues the functional deficit observed in Bag3 mutants

As Amiodarone and Metformin significantly ameliorate the protein aggregate aspect of myofibrillar myopathy in zebrafish or zebrafish and human myoblasts respectively, we wished to further examine their potential to prevent fibre disintegration and improve muscle function, which is ultimately the desired outcome. As such, 5-somite staged *bag3*^−/−^ mutant embryos were dechorionated and incubated in unsupplemented E3 embryo medium or in E3 medium containing Metformin or Amiodarone. Following a 12 hour treatment, the embryos were placed in methyl cellulose, with or without the drug, for 2 hours to promote fibre disintegration, and subsequently fixed and labelled to examine fibre integrity (Figure 8A). Whilst there was a significant increase in the number of phenotypic embryos following Metformin or Amiodarone treatment (Figure 8B), both drugs resulted in a striking reduction in the frequency of fibre disintegration, when compared to untreated *bag3*^−/−^ embryos (Figure 8A & 8C).

**Figure 8:**
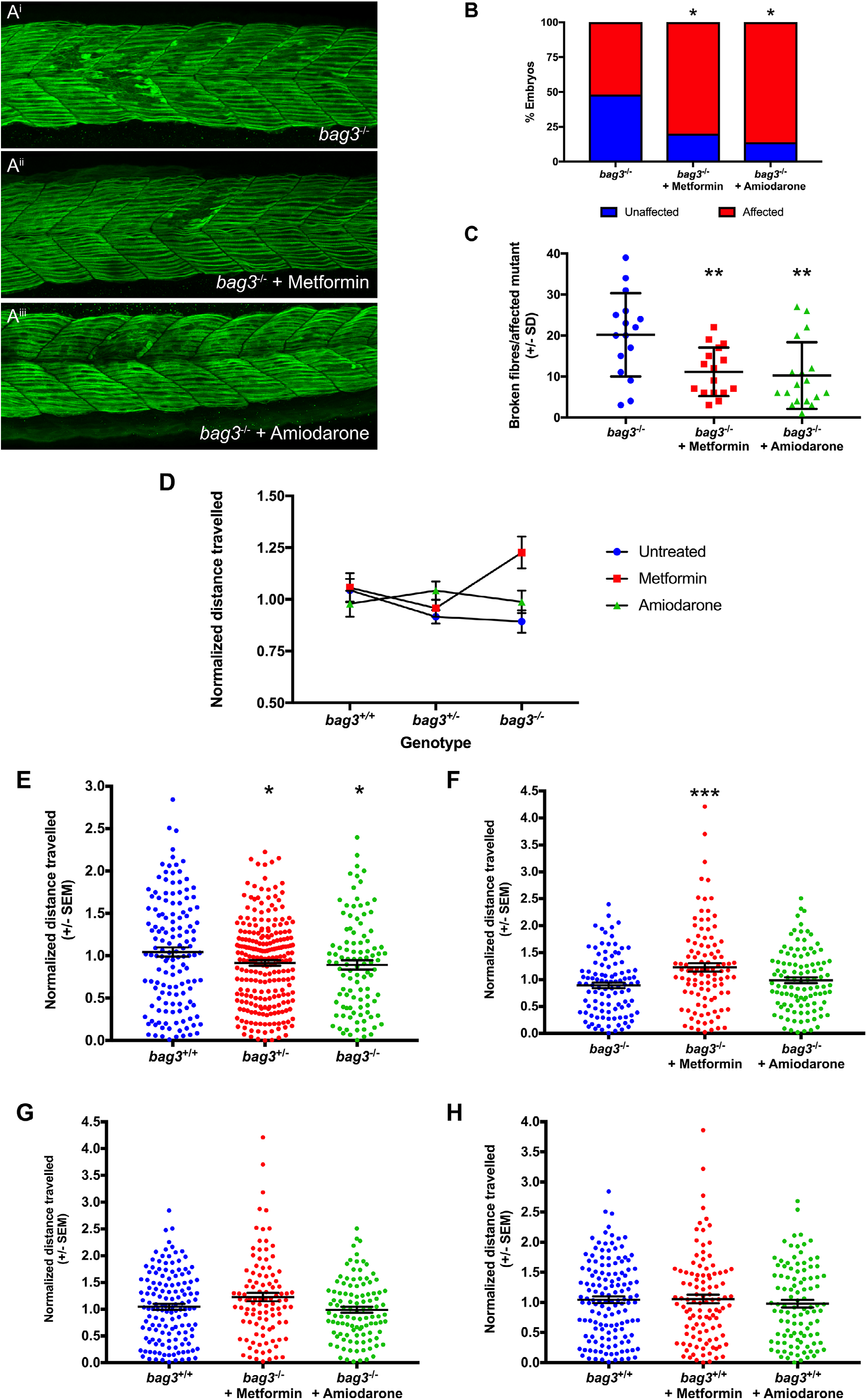
Metformin prevents fibre disintegration and improves muscle function in Bag3 mutant embryos. (A) Representative maximum projection images of Myosin labelled 26 hpf; untreated, Metformin treated, and Amiodarone treated embryos following incubation in methyl cellulose. (B) Metformin or Amiodarone treatment results in an increase in the proportion of *bag3*^−/−^ embryos that display damaged muscle fibres. (C) Quantification of disintegrated fibres in vehicle-, Metformin-, and Amiodarone-treated *bag3*^−/−^ embryos fish reveals that both drug treatments result in a significant reduction. (D) Analysis of distance travelled by two-way ANOVA identifies a significant interaction between genotype and drug treatment (p=0.005). (E) Both *bag3*^+/−^ and *bag3*^−/−^ fish show a significant reduction in distance travelled compared to *bag3*^+/+^ fish. (F) Metformin, but not Amiodarone, significantly increases the distance swum by *bag3*^−/−^ fish, compared to untreated *bag3*^−/−^ controls. (F) The distance swum by Metformin treated *bag3*^−/−^ fish is comparable to that of untreated *bag*^+/+^ controls (F) The distance swum by Metformin or Amiodarone treated *bag3*^+/+^ fish is not significantly different to that of untreated *bag3*^+/+^ controls. (G) Treatment with either Metformin or Amiodarone has no effect on *bag3*^+/+^ fish. Error bars represent +/-SEM with four-six independent biological replicates. The number of fish in each replicate is documented in Supplementary Table 5. *p<0.05, **p<0.01, ***p<0.001, ****p<0.0001 calculated using a Chi-square test or a one way ANOVA with Dunnett’s post-hoc multiple comparison correction test.

To confirm that this reduction was due to an increase in autophagy, we examined LC3 accumulation in 28 hpf untreated, NH_4_Cl treated, Metformin & NH_4_Cl treated, and Amiodarone & NH_4_Cl treated wildtype embryos. We observed an increase in LC3 accumulation in both Metformin & NH_4_Cl treated and Amiodarone & NH_4_Cl treated wildtype embryos, compared to embryos treated with NH_4_Cl alone (Supplementary Figure 4A), demonstrating that a 14 hour treatment with Metformin or Amiodarone is sufficient to upregulate autophagy *in vivo*, and subsequently reduce fibre disintegration.

We further validated the therapeutic potential of Metformin and Amiodarone by examining their capacity to improve muscle function. Our initial studies revealed that the drug doses used at the 36 hpf, and 5-somite stages were toxic at 6 dpf. We therefore conducted a drug toxicity assay to determine a well tolerated, non toxic dose, to use on 6 dpf larvae for our functional assays. We tested the effect of 5 concentrations of Metformin and Amiodarone on fish survival and swimming performance. Whilst larvae treated with the highest conccentration of Metformin (10 μM) showed no signs of toxicity, an equivalent 10μM dose of Amiodarone resulted in larval death. Treatment with 1μM of Amiodarone had no toxic effects and as such this concentration was used for subsequent experiments (Supplementary Figure 6).

Having identified suitable non-toxic levels we treated 5 dpf old *bag3*^−/−^ with 1 μM Amiodarone, 10 μM Metfromin, or unsupplemented E3 embryo medium, for 16 hours after which the locomotion assays were performed. A two way ANOVA revealed a significant interaction between genotype and drug treatment (p=0.005; Figure 8D, Supplementary Table 5). We found that as per our previous experiments *bag3*^−/−^ mutants had a significant reduction in distance swum (Figure 8E). Whilst Amiodarone treatment had no significant effect on muscle function of *bag3*^−/−^ larvae, Metformin treatment dramatically improved the distance swum by *bag3*^−/−^ fish compared to untreated *bag3*^−/−^ fish (Figure 8F). Remarkably, the mean distance swum by Metformin treated *bag3*^−/−^ larvae was no longer different to untreated *bag3*^+/+^ larvae indicating that Metformin completely restored muscle function in the mutant fish (Figure 8G). To examine if this increased swimming performance in Metformin treated *bag3*^−/−^ larvae was due to a general increase in motility of the fish, irrespective of the *bag3* mutation, we compared the locomotion of untreated and Metformin treated *bag3*^+/+^ larvae. Treatment of *bag3*^+/+^ with Metformin had no effect on their swimming distance, demonstrating that the improvement in muscle function seen in *bag3*^−/−^ is specific, and not a generalized response (Figure 8H). We also assessed the capacity of Metformin and Amiodarone to increase autophagic flux *in vivo* at 6 dpf. Whilst Amiodarone had no effect on LC3 accumulation, Metformin resulted in LC3 accumulation that was higher than that seen in embryos treated with NH_4_Cl alone (Supplementary Figure 5B), highlighting that at the doses used for the swimming assays Metformin, but not Amiodarone, stimulates autophagy explaining the increased muscle function in *bag3*^−/−^ larvae.

## Discussion

In the current study we have generated novel zebrafish models of BAG3 myofibrillar myopathy, which recapitulate the hallmark features of the disease. Our data collectively demonstrates that the FDA-approved, autophagy stimulating drug, Metformin reduces protein aggregation, prevents fibre disintegration, and rescues muscle weakness in our model, making it a very promising therapy for the treatment of BAG3 myofibrillar myopathy, and the wider group of myofibrillar myopathies.

Using the models we have generated, we reveal that the overexpression of BAG3^P209L^ results in protein aggregation, the majority of BAG3^P209L^ being in the insoluble fraction. Despute this, BAG3^P209L^ is functional and can rescue the loss of function. This is consistent with the proposal that the formation of protein aggregates results in sequestration of functional BAG3, both BAG3^P209L^ and BAG3^wt^, ultimately causing insufficiency ^26^ and analyses showing BAG3^P209L^ is correctly folded and functional ^27^. Therefore, whilst the aggregation of BAG3^P209L^ is the initial trigger for the disease, it does not directly cause muscle weakness. The loss of Bag3 on the other hand, results in fibre disintegration and impaired muscle function, due to reduced autophagic activity. We further confirm that this reduced autophagic activity is evident in skeletal muscle from BAG3 MFM patients, consistent with previous reports of defective autophagy BAG3 cardiomyopathy ^28^. The BAG3^P209L^ expressing fish therefore provides a model for the early stages of BAG3 myofibrillar myopathy, where aggregates are present but muscle performance is unaffected, and the loss of function Bag3 mutant provides a model for the later stages of the disease in which, fibre disintegration and muscle weakness is evident.

Given that the presence of aggregates triggers the loss of Bag3 and associated muscle weakness, we examined autophagy up regulation as a strategy to promote aggregate clearance. We tested 71 compounds all of which have been previously shown to stimulate autophagy. Despite this, only 13% of compounds significantly reduced protein aggregates. Whilst this relatively low number of successful hits may be attributed to the use of all compounds at a single concentration of 10 μm, it may also be explained by the selective nature of autophagy stimulating drugs, and their mechanism of action. For example, Lithium ^29^, Rilmenedine ^30^, and Trehalose ^31,32^ have previously been shown to reduce protein aggregation in models of neurodegenerative diseases, although they have failed to show beneficial effects in patients, potentially due to difficulties in achieving therapeutic levels. However, in the current study none of these drugs had an effect on the number of BAG3^P209L^ aggregates, suggesting that autophagy stimulating drugs may be selective in the type of aggregates they target and/or the tissue in which they act. There is therefore a need for specific models to evaluate therapies for each of the protein aggregate disorders.

Of the 9 drugs that successfully reduced protein aggregates in the zebrafish BAG3^P209L^ expressing fish, two compounds, Amiodarone and Metformin, are currently approved for clinical use by the US Food and Drug Administration board. Amiodarone is known to induce neuropathy in some cases and, even more, rarely myopathy ^33,34^ As a result Amiodarone would not be indicated for treating BAG3 myopathy. Metformin on the other hand, is the most effective compound we identified. It is the most widely prescribed drug for the treatment of diabetes, with extensive documentation on its tolerability and safety and an outstanding risk to benefit profile. Metformin has been shown to exert its therapeutic benefits through several different pathways including AMP protein kinase dependent and independent mechanisms, modulating mitochondrial oxidative phosphorylation (reviewed in ^35^), and most recently by altering the gut microbiome ^36^. Whilst we believe the increase in autophagy flux following Metformin treatment is responsible for the improvement in muscle pathology and function seen in our BAG3 myofibrillar myopathy models, the modulation of these other pathways may also be contributing.

Remarkably, in addition to clearing protein aggregates, Metformin prevents the fibre disintegration and muscle weakness seen in the loss of function Bag3 mutant. Metformin treatment in BAG3 myofibrillar myopathy patients would therefore have dual benefits: increasing autophagy, promoting aggregate clearance and preventing loss of functional BAG3 by sequestration; additionally, the increase in autophagy levels would compensate for the reduced autophagic levels due to reduction in BAG3. Metformin therefore has potential to address both of the muscle phenotypes associated with the disease and is readily applicable to the treatment of BAG3 myofibrillar myopathy. In addition, given the conserved impairment in autophagy, Metformin may also be suitable for the treatment of cardiomyopathy resulting from mutations in BAG3, and other myofibrillar and aggregate myopathies.

## Methods

### Fish maintenance

Fish maintenance and handling were carried out as per standard operating procedures approved by the Monash Animal Services Ethics Committee. The generation of transgenic and mutants strains was approved by the School of Biological Sciences Animal Ethics Committee (BSCI/2013/27 and BSCI/2015/06 respectively). All experiments were carried out on embryos of TU/TL background. Fish were anesthetized using Tricaine methanesulfonate (3-amino benzoic acidethylester, Sigma) at a final concentration of 0.16% in E3 embryo medium (5 mM NaCl, 0.17 mM KCl, 0.33 mM CaCl, 0.33 mM MgSO4 in water).

### Human Ethics

This study was performed according to the guidelines of the Committee on the Use of Human Subjects in Research of the Policlinico Hospital of Milan (Milan, Italy) or the Institute of Myology, Pitié-Salpêtriere hospital. Informed consent was obtained from all family members.

### Generation of transgenic and mutant strains

Transgenic constructs were assembled using the multisite gateway cloning kit ^37^ with β-actin promoter in the 5’ entry position, loxP-mCherry-pA-loxP (Genbank accession: KF753698) in the middle entry position, and C-terminal, eGFP tagged, human full length wildtype BAG3 or BAG^P209L^ ^26^ in the 3’ entry position. The constructs were injected at 25 ng/μl into the one-cell-stage embryos along with transposase RNA (25ng/μl) that was synthesized from the pcs2FA-transposase vector ^37^using the mMessage machine Sp6 kit (Ambion). Transgenic strains generated were *Tg(βAct:loxP-mCherry-pA-loxP:Hs.BAG3^wt^-eGFP), Tg(βAct:loxP-mCherry-pA-loxP:Hs.BAG3^P209L^-eGFP)*. Crossing of either of these two strains to the *Tg(actc1b:iCre)*^38^ strain results in the excision of the loxP-mCherry-pA-loxP cassette and subsequent generation of *Tg(βAct:Hs.BAG3^wt^-eGFP)* and *Tg(βAct:Hs.BAG3^209L^-eGFP)* hereafter referred to as Tg(BAG3^wt^-eGFP) and Tg(BAG3^P209L^-eGFP) respectively. The BAG3 mutant strain was generated using CRISPR/Cas9 technology as per ^36^ with the following amendments: the guide RNA was synthesized according to ^39^. The guide RNA was injected at 120 ng/μl into the one-cell-stage embryos, along with Cas9 protein and Cascade Blue dye (Molecular Probes). All primers used for the generation, and genotyping of the BAG3 mutant strain are listed in supplementary Table 1.

### Immunofluorescence and western blot experiments

Zebrafish immunofluorescence experiments were performed accordingly to previously published protocols ^40^. Primary antibodies used in this study were anti-eGFP (Invitrogen, A-11122, 1:150) and anti-actinin2 (Sigma, A7811, 1:100). Stained embryos were mounted in 1% low melting point agarose and imaged using the Zeiss LSM 710 confocal microscope. The maximum intensity projections were obtained using Fiji (http://fiji.sc).

For zebrafish western blot assays protein lysates were obtained as per ^41^ and quantified using the Qubit fluorometric quantification (Thermo Fisher Scientific). 30 μg of each sample, along with reducing agent (Life Technologies) and protein loading dye (Life Technologies), was heated at 70°C for 10 min, separated by SDS-PAGE on NuPAGE 4–12% Bis-Tris gels, and transferred onto PVDF membrane (Millipore). Following transfer, the membrane was blocked with 5% skimmed milk in PBST and subsequently probed with anti-LC3 (Cell Signaling, 12741, 1/2000), washed and incubated with HRP-conjugated secondary antibody (1:10 000, Southern Biotech). Immunoblots were developed using ECL prime (GE healthcare) and imaged using a chemiluminescence detector (Vilber Lourmat). The membrane was subsequently stripped by incubating in 1X stripping buffer (200mM glycine, 0.1% SDS, 1%Tween20, pH 2.2) twice for 10 minutes, washed in PBST, and stained with Direct blue as per ^42^ to detect total protein. The blot images were quantified using Image Lab software (Bio-rad) and a one-way ANOVA statistical test was used to test for significant changes in LC3 accumulation.

Details on human muscle biopsies are provided in Supplemetary Table 7. For immunohistochemistry, sections were re-cut from stored frozen muscle (–80 °C). 8 μm thick cryo-sections were air-dried and fixed in cold acetone at –20 °C for 10 min. Staining was carried out using an automated immunostainer (BenchMark XT, Ventana medical systems) as per manufacturers instructions with anti-p62 antibody (BD Biosciences). For western blot the biopsy samples were isolated from the patient and homogenized in a lysis buffer containing 20mM Tris-HCl (pH 7.8), 140mM NaCl, 1mM EDTA, 0.5% NP40, 1mM phenylmethylsulfonil fluoride, and complete protease inhibitor mixture (Roche Diagnostics, Rotkreuz, Schweiz), with a POTTER S Homogenizer (B.Braun Biotech International-Sartorius group). Samples were pulsed 5 times for 5 seconds each at a speed of 1000rpm. Samples were then passed 5 times through a 30.5-gauge needle to disrupt the nuclei, then incubated at 4°C for 15 min and finally centrifuged at 13000 rpm for 15 min at 4°C. Total protein concentration was determined according to Bradford’s method. Samples were resolved on 12% polyacrylamide gel for LC3B (1:1000, Cell Signaling Technology, USA) and SQSTM1/p62 (1:1000, Millipore, Temecula, CA, USA) antibodies and transferred to nitrocellulose membranes (Bio-Rad Laboratories, Hercules, CA, USA). We also determined the expression for each sample of anti-GAPDH (1:600) (SIGMA, Saint Louis, MO, USA) housekeeping protein. Detection was performed with anti-rabbit and anti-mouse horseradish peroxidase (HRP)-conjugated secondary antibodies respectively (DakoCytomation, Carpinteria, CA, USA), followed by enhanced chemiluminescence (ECL) development (Amersham Biosciences, Piscataway, NJ, USA). Bands were visualized by Odissey system. Each sample was evaluated in three independent experiments.

### Drug screen and automated aggregate quantification

The SCREEN-WELL Autophagy library (ENZO Life Sciences) containing 71-autophagy inducing compounds was used in this study. 32 hpf Tg(BAG3^P209L^-eGFP) embryos were placed in wells of a 24-well plate (5 embryos/well) following which drug was added to a final concentration of 10 μM in 1ml volume. E3 embryo media without any drug supplementation, and E3 embryo media containing 0.1% DMSO were used as controls. Embryos were incubated in their respective drugs (or control solutions) for 16 hours following which, fixed in 4% paraformaldehyde for 2 hours, washed into PBST and mounted in 1% low melting point agarose for imaging using the Zeiss LSM 710 confocal microscope.

A semi-automated workflow for detecting aggregates in three-dimensional confocal image stacks was developed using ImageJ and the ImgLib2 library ^43,44^ Briefly, a median filter with a radius of one pixel was applied to the image volume to reduce overall shot noise. To calculate overall fish muscle tissue volume, image stacks were smoothed further by applying a Gaussian filter with a radius of five pixels and then segmented using an Otsu automatic threshold ^45^. The fish muscle tissue was delineated as the largest connected component in the field of view resulting in exclusion of small spurious segmented particles. Aggregates were enhanced by applying a Difference of Gaussian (DoG) blob enhancing filter to the median filtered image volume, where the estimated radius of the blob (i.e., aggregate) was set at 1um. Aggregates were then detected in the DoG filtered image volume using a local neighbourhood maxima detection algorithm with a minimum peak intensity cut-off of 2,500 (in a 12-bit greyscale intensity range). Detected aggregates were masked using the segmented muscle tissue region to select only the aggregates within the tissue. Aggregate density was calculated as the number of aggregates per unit volume of tissue. Source code and a current build for the ImageJ/Fiji Aggregate plugin can be found at: https://gitlab.erc.monash.edu.au/mmi/aggregate. All drugs were tested in triplicate with the treatment and imaging order randomized within each replicate, and the identity of treatment blinded.

### Movement assays

Touch-evoked response assay to examine maximum acceleration and assays to determine distance swum were performed on 2 dpf embryos and 6 dpf larvae respectively, as per ^38,46^. For experiments on the effect of drug treatment on distance swum, larvae were placed in individual wells of a 24 well plate following which the epMotion liquid handling robot (Eppendorf) was used to automatically dispense drugs into individual wells to a total volume of 1ml, in a randomized order. The fish were incubated in their respective drugs for 14-16 hours after which the locomotor activity described above were performed.

### Electron Microscopy

Zebrafish were fixed according standard procedures in 2.5% glutaraldehyde, 2% paraformaldehyde in 0.1M sodium cacodylate buffer. Post-fixed with 1% OsO4, 1.5% K3Fe(III)(CN)6. Dehydration was done with ethanol and the zebrafish were flat embedded in Epon 812. Ultrathin sections of 70nm were cut on a Leica Ultracut UCT7 and stained with uranyl acetate and lead citrate. Large area EM tile sets were taken on a FEI NovaNanoSEM 450 equipped with an ETD secondary electrons in-lens detector set at 10kV and a STEM II (HAADF) detector set at 30kV. MAPS 2.1 software was used to create the tile sets. High resolution EM imaging was done on a Hitachi 7500 TEM and a FEI Tecnai 12 TEM.

### Drug concentrations

For the initial drug screen and fibre disintegration rescue experiments all compounds were tested at 10 uM. For functional validation of the top FDA approved compounds a dose-toxicity assay identified optimal concentrations of 10 uM for Metformin and 1 uM for Amiodarone. To examine autophagic flux in early drug treated embryos (fibre disintegration rescue experiments) ammonium chloride was used at 100 uM for 4 hours. To examine autophagic flux in 6 dpf *bag3* mutants and in drug treated larvae (locomotion rescue experiments) ammonium chloride was used at 100 uM for 12-14 hours.

### Cell culture

Anonymous immortalized healthy control myoblast lines were provided by Eric A. Shoubridge (Department of Human Genetics, McGill University, Quebec, Canada). Briefly, human myoblasts were cultured from biopsy material and transduced with retroviral vector expressing HPV16-E6/E7 and with another one expressing the catalytic component of human telomerase (htert), to increase their lifespan in vitro ^47,48^. The immortalized human myoblasts were cultivated in growth medium (GM, SkMAX skeletal muscle medium with supplements (Wisent, Kit #301-061-CL) containing 20% heat inactivated fetal bovine serum) in a humidified incubator at 37°C, 5% CO_2_. Cells were passaged using 0.05% trypsin when they have reached 70-80% confluency (about every 3-4 days) and kept in culture not longer than passage 15.

### Differentiation and transduction of immortalized human myoblasts

Immortalized human myoblasts were plated at a density of 1×10^6^ cells /35mm dish on plastic in 2 ml growth medium. To induce differentiation into myotubes, cells were washed once with 1xPBS, and differentiation medium (DM, DMEM containing 2% Horse Serum) was added for the indicated time of differentiation. At day2 (D2), cells were transduced with Ad-BAG3-GFP wild type or Ad-BAG3(P209L)-GFP, or with 1 PFU of Ad-GFP only as control, as described ^49^. Briefly, myoblasts were incubated with adenoviruses in 400μl of medium for 1h before adding 1.6 ml medium. The next day, cells were washed once with 1x PBS and 2 ml of DM was added. Different batches of Horse Serum have been tested for transduction efficiency.

### Myoblast antibodies and chemicals

The following antibodies and drugs were used: rabbit anti-BAG3 LP10 was raised against a C-terminal peptide (SSMTDTPGNPAAP) of human BAG3 fused with glutathione S ^16^; Metformin hydrochloride (ab120847, Abcam) and Amiodarone (ab141444, Abcam) were used at 100 μM and 15 μM respectively for 48 hours; Hoechst Bisbenzimide H33342 (B2261, Sigma).

### Myoblast immunofluorescence and immunoblotting

Immortalized human myoblasts were washed twice with 1 ml Luftig buffer (0.2M sucrose, 35mM PIPES, pH 7.4, 5mM EGTA, 5mM MgSO4) and fixed in 4% paraformaldehyde in PBS for 25 min at 37°C. DNA was stained with a cell permeable Hoechst in PBS for 15 min at room temperature. Specimen were washed 2x with PBS-M (1xPBS containing 1mM MgCl_2_) and post-fixed in PBS+3.7% formaldehyde for 20 min at 37°C ^49^. Cells were subsequently washed twice with PBS-M, once with H_2_O and mounted. Epifluorescence images were acquired with an AxioObserver Z1 system using a 40x Plan-Neofluoar 0.6NA objective and a charged-coupled device (CCD) camera Axiocam MRm controlled by the Zen software (Carl Zeiss). Confocal microscopy of fixed cells was performed with a Perkin Elmer UltraVIEW Spinning Disc Confocal (40x 0.75NA), equipped with an EMCCD cooled charge-coupled camera at −50°C (Hamamatsu Photonics K.K) and driven by Volocity software version 6.01.

The volocity software version 6.0 (Quorum Technologies) and Image J 1.48v (National Institute of Health) software were used for processing on entire images before cropping to emphasize the main point of the image. Processing was limited to background subtraction, brightness/contrast adjustment and deconvolution.

For immunoblotting, equal amounts of proteins were loaded on SDS-PAGE and analyzed by Western blot as described before ^50^. Protein concentrations were determined using the DC^M^ Protein Assay Kit BioRad (#500-0116, Biorad).

### Triton soluble and insoluble cell fractionation

Immortalized human myoblasts were washed once with 2 ml pre-warmed 1xPBS and detached with 1 ml pre-warmed PBS-Citrate-EDTA (13.6 mM Sodium citrate, 0.6 mM EDTA pH8 in 1xPBS) for 3-5 min in a humidified incubator at 37°C, 5% CO_2_. Cells were carefully transferred into a 1.5 ml plastic tube and centrifuged for 5 min at 1000 rpm at 4°C. The supernatant was removed and the pelleted cells lysed by 3-cycles of freezing/thawing in 150 μl lysis buffer (20 mM Tris-HCl pH 7.6, 150 mM NaCl, 1 mM EDTA, 0.1% IGEPAL, 1 mM DTT, 1x Complete Protease Inhibitor [Roche]). Cell extracts were centrifuged for 5 min at 12000 rpm, 4°C and the triton soluble fraction was collected in SDS sample buffer (62.5 mM Tris-HCl pH 6.8, 2.3 % SDS, 10 % glycerol, 5 % β-mercaptoethanol, 0.005 % bromophenol blue, 1 mM phenylmethylsulfonyl fluoride). The triton insoluble pellet was washed 1x with 150 μl lysis buffer, centrifuged for 5 min at 12000 rpm at 4°C and resuspended in SDS sample buffer. After aspirating the supernatant, the pellet was resuspended in 150 μl lysis buffer and 75 μl 3xTEX was added. Protein quantification was performed on the soluble fraction and equal amounts of soluble and insoluble fractions were loaded on SDS-PAGE and analyzed by western blot.

### Analysis of aggregate intensity and volume

Z-stacks of fixed cells were acquired using a Perkin Elmer UltraVIEW Spinning Disc Confocal (40x 0.75NA). The background was removed for each acquisition before cropping representative cells for each treatment by drawing semi-automatically a line around the selected cells. The P209L aggregates volume and intensity were measured on the complete Z-stack using the velocity software. The ratio between volume/intensity of aggregates versus the complete cell was quantified.

### Analysis of aggregates versus diffuse staining

Epifluorescence images of 40 randomly chosen fields per condition were acquired with an AxioObserver Z1 system using a 40x Plan-Neofluoar 0.6NA objective. For each field, the percentage of cells or myotubes containing aggregates and/or filaments was counted semi-automatically using the Image J 1.48v software. Statistical calculations were performed using Prism 6.0 (GraphPad Software) statistical software. The statistical tests used are indicated in the figure legends.

### Statistical analysis

GraphPad Prism and SPSS statistics packages were used to analyse data in this study. Note that the number of independent biological replicates examined for each experiment and the number of fish used within each replicate combined with the significance tests used and the associated t/F value, degrees of freedom (df) and exact p values obtained are detailed in Supplementary Tables 2 - 6.

#### Drug screen

To identify drugs that significantly reduced protein aggregation in Tg(BAG3^P209L^-eGFP) embryos a one way ANOVA statistical analysis was performed with treatment as a fixed effect and replicate as a random effect, and Dunnett’s post-hoc test was used to correct for multiple comparisons.

#### Movement assays

Outliers, identified as being more than 1.5 times the interquartile range below the 1^st^ quartile or above the 3^rd^ quartile in each treatment group were omitted. Individual data points were normalized to the mean of all data points from that replicate, following which a t-test or a one way ANOVA with Dunnett’s post-hoc multiple comparison correction test was performed to identify significant differences. For experiments examining the ability of the drugs to rescue the movement deficit observed in the mutant, a two way ANOVA was performed. Having shown a significant interaction between drug and genotype, one way ANOVAs with Dunnett’s post-hoc multiple comparison correction test were performed to explore the basis of the interaction.

#### Fibre disintegration rescue experiments

Outliers were identified using the Prism Rout (Q=1%) method and omitted. Chi-square test was used to compare the change in the number of phenotypic fish following drug treatment. A one way ANOVA with Dunnett’s post-hoc multiple comparison correction test was used to compare the number of broken fibres/phenotypic fish.

#### Autophagic flux assays

The normalized change in LC3 levels between the different genotypes were compared using a one way ANOVA with Dunnett’s post-hoc multiple comparison correction test.

### Data availability

Raw data and files for statistical analysis are available at figshare and can be accessed using the following link: https://monash.figshare.com/projects/Metformin_rescues_muscle_function_in_BAG3_myofibrillar_myopathy_models/60959.

## Acknowledgements

We would like to thank Prof Bernard Brais for the myoblast cell lines utilised. Drs Aidan Sudbury and Keyne Monro for statistical advice, and the patients who contributed samples to this study. We would also like to thank the Bellini Foundation and Fondazione Roby for funding this work.

