## Supplementary Figures and Tables for "Metformin rescues muscle function in BAG3 myofibrillar myopathy models"

Avnika A. Ruparelia[1], Emily A. McKaige[1], Caitlin Williams[1], Keith E. Schulze[2], Margit Fuchs[3,4], Viola Oorschot[5], Emmanuelle Lacene[6], Meregalli Mirella[7], Emily C. Baxter[1], Yvan Torrente[7], Georg Ramm[5,8], Tanya Stojkovic[9], Josée N. Lavoie[3,4,10], and Robert J. Bryson-Richardson[1*]

[1] *School of Biological Sciences, Monash University, Melbourne 3800, Australia*

[2] *Monash Micro Imaging, Monash University, Melbourne 3800, Australia*

[3] *Centre de Recherche sur le Cancer de l'Université Laval, Ville de Québec, Quebec, Canada.*

[4] *Oncologie, Centre de Recherche du Centre Hospitalier Universitaire (CHU) de Québec-Université Laval, Ville de Québec, Quebec, Canada.*

[5] *Monash Ramaciotti Centre for Structural Cryo-Electron Microscopy, Monash University, Melbourne 3800, Australia*

[6] APHP, Centre de Référence de Pathologie Neuromusculaire Nord/Est/Ile-de-France, Institut de Myologie, Laboratoire de Pathologie Risler, GH Pitié-Salpêtrière, Paris, France.

[7] *Stem Cell Laboratory, Department of Pathophysiology and Transplantation, Università degli Studi di Milano, Fondazione IRCCS Ca' Granda Ospedale Maggiore Policlinico di Milano, Centro Dino Ferrari, via F Sforza, 35, 20122 Milan, Italy*

[8] *Biochemistry and Molecular Biology, Monash Biomedicine Discovery Institute, Monash University, Melbourne, Victoria, Australia.*

[9] *Centre de référence des maladies neuromusculaires, Hôpital Pitié-Salpétrière, Assistance-Publique Hôpitaux de Paris, Paris, France.*

[10] *Département de Biologie Moléculaire, Biochimie Médicale et Pathologie, Université Laval, Ville de Québec, Quebec, Canada.*

[*]Corresponding author.

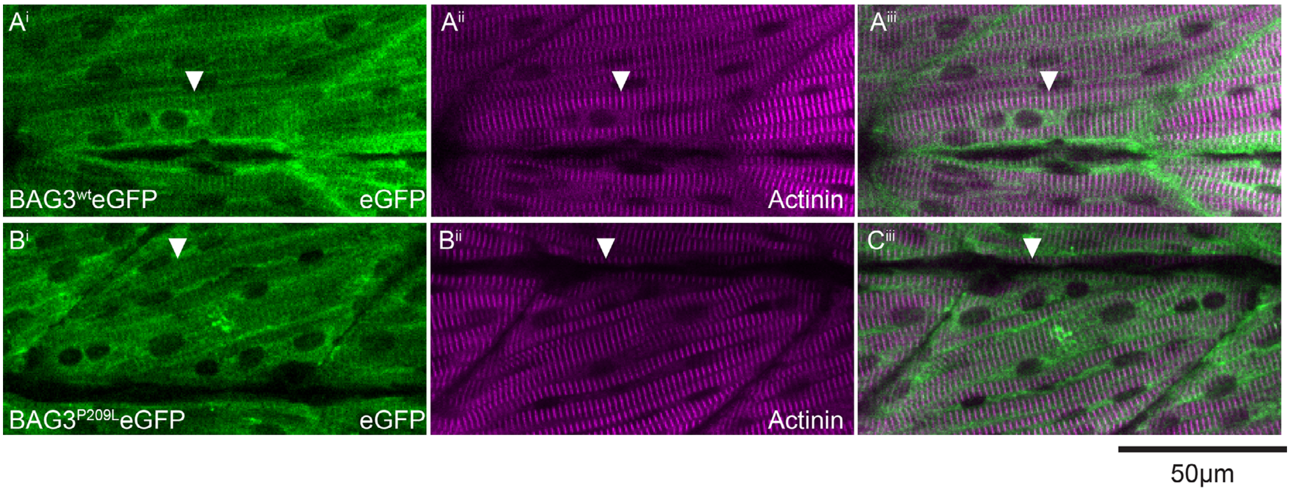


**Supplementary Figure 1**: Z-disk localization of $BAG3^{wt}$–eGFP (A) and $BAG3^{P209L}$–eGFP (B) as shown by the overlap between GFP (green) with Actinin (red) staining (arrowhead).

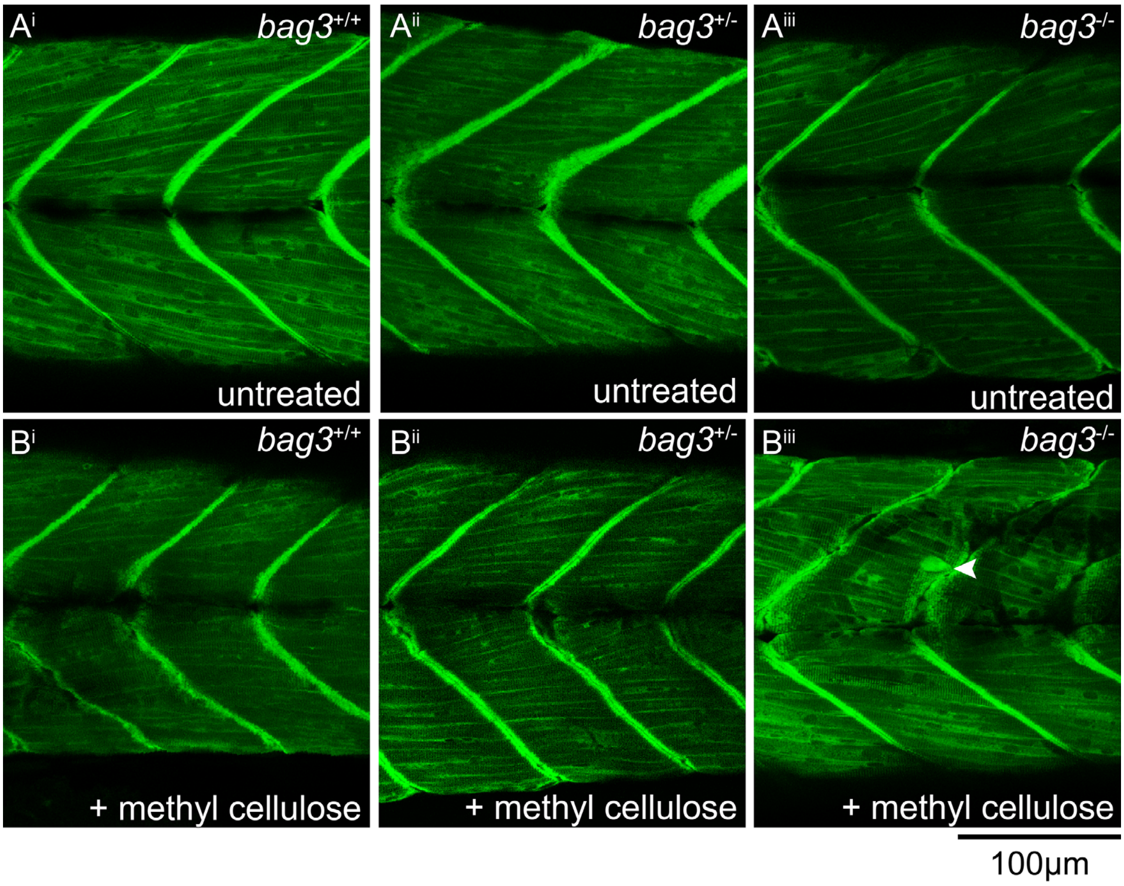


**Supplementary Figure 2:** Bag3 mutants have normal, intact muscle fibres at 6-dpf. (A) Live confocal images of *bag3*$^{+/+}$, *bag3*$^{+/-}$, and *bag3*$^{-/-}$ on a Tg(FLNC-eGFP) background showing normal muscle structure in all there strains (B) Incubation of *bag3*$^{+/+}$, *bag3*$^{+/-}$, and *bag3*$^{-/-}$ embryos expressing FLNC-eGFP in the muscle in methyl cellulose induces mild fibre disintegration in *bag3*$^{-/-}$ embryos.

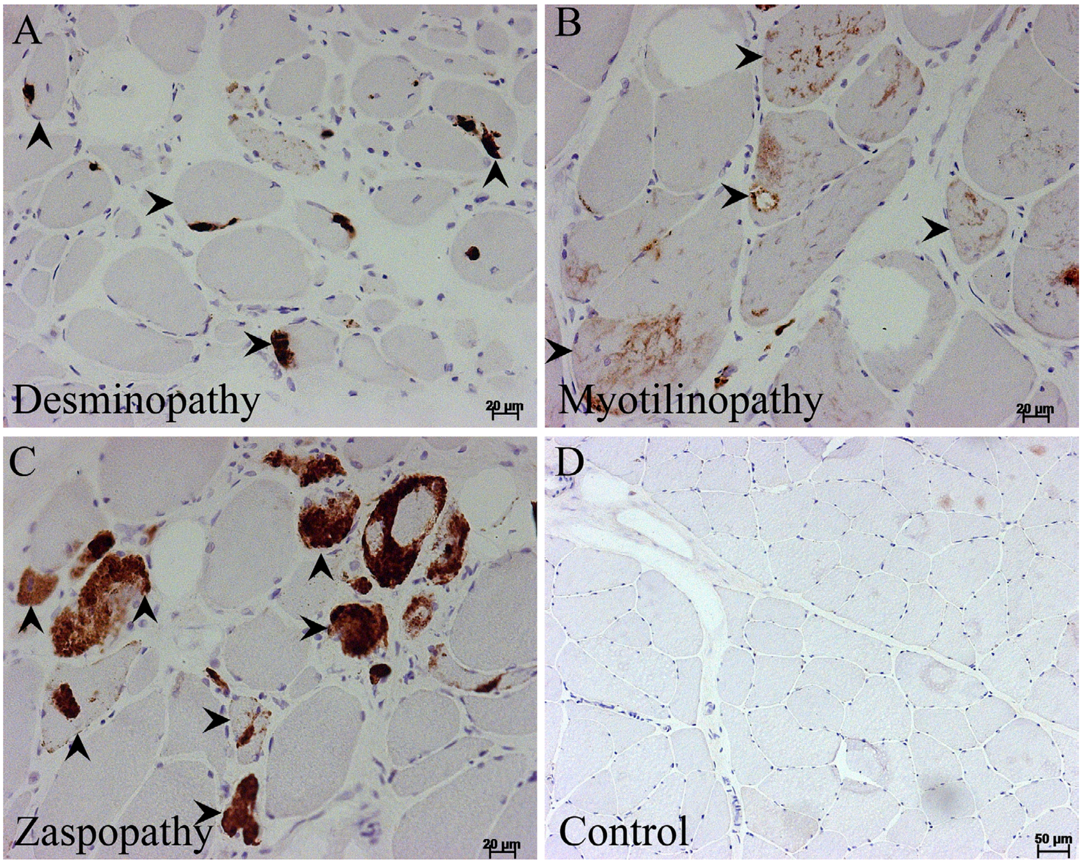


**Supplementary Figure 3:** Impaired autophagy, evidenced by abnormal accumulation of p62, is a conserved feature seen in myofibrillar myopathy caused by mutations in Desmin (A), Myotilin (B), and ZASP (C). Control samples show no evidence of p62 accumulation (D).

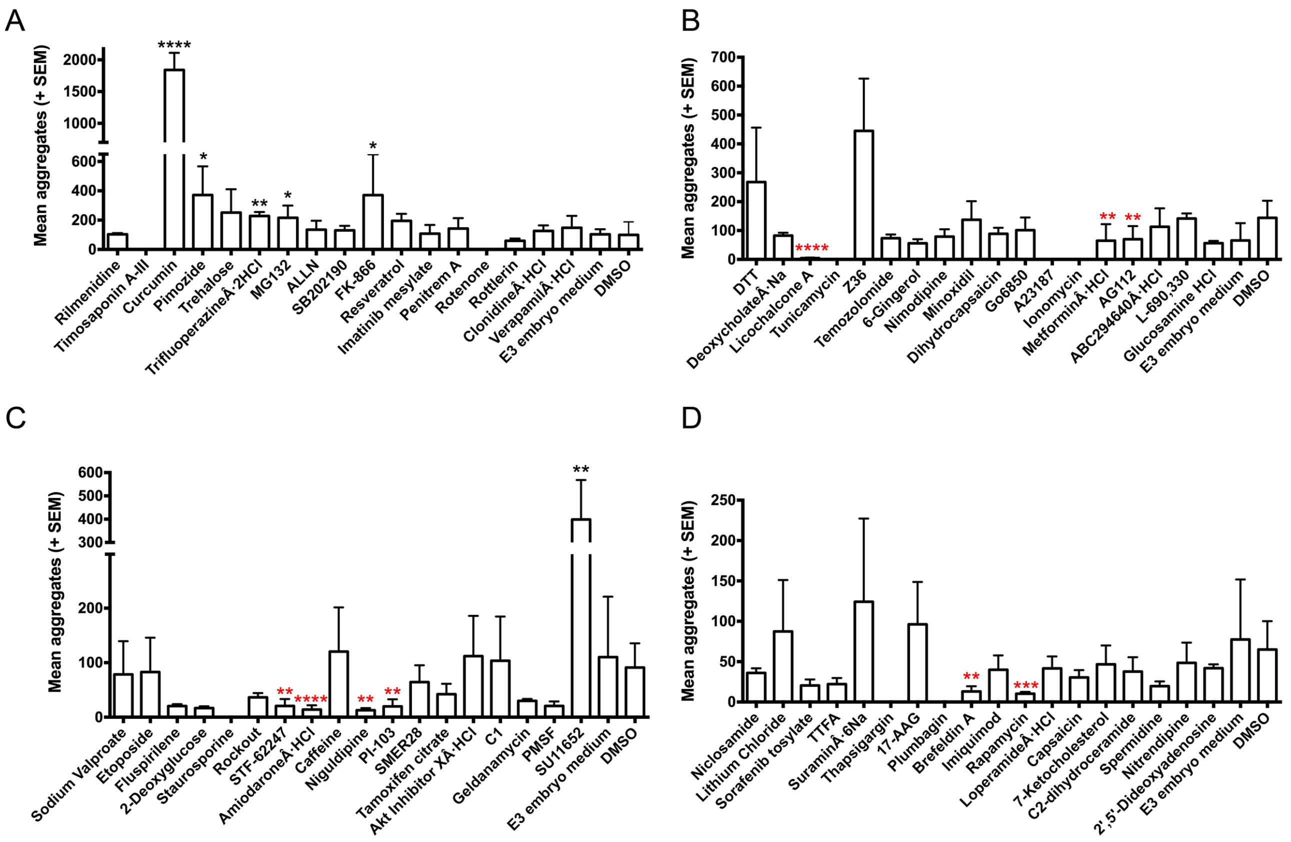


**Supplementary Figure 4**: Evaluation of autophagy stimulation in promoting aggregate clearance. For the purposes of feasibility, the 71 autophagy stimulating compounds were randomly allocated to one of four groups with each group having its own E3 embryo medium and DMSO control treatment. The mean number of aggregates for the various compounds in each group is displayed in graphs A-D. Error bars represent SEM with three independent replicates. The number of fish in each replicate is documented in Supplementary Table 4. Asterisks in black show drugs that significantly increased the number of aggregates compared to DMSO control, and asterisks in red mark the compounds that significantly reduced protein aggregates compared to DMSO control. *p<0.05, **p<0.01, ***p<0.001, ****p<0.0001 calculated using a one way ANOVA with Dunnett’s post-hoc multiple comparison correction test.

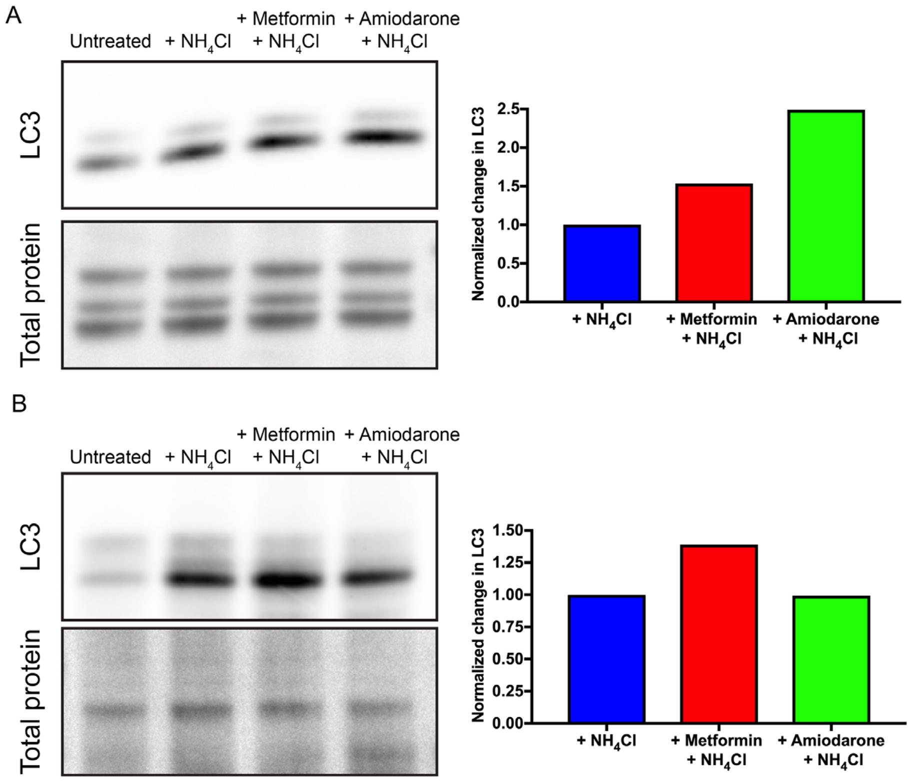


**Supplementary Figure 5:** Examination of autophagic flux in drug treated embryos. Western blot for LC3 and Direct Blue stain for total protein (loading control) on protein lysates obtained from 28 hpf (A) or 6 dpf (B) wildtype fish that were untreated, ammonium chloride ($NH_4Cl$) treated, Metformin and $NH_4Cl$ treated or Amiodarone and $NH_4Cl$ treated fish. The autophagic flux determined by normalizing LC3 levels to that of total protein. The change in LC3 levels following $NH_4Cl$ treatment is presented relative to the levels in the untreated groups.

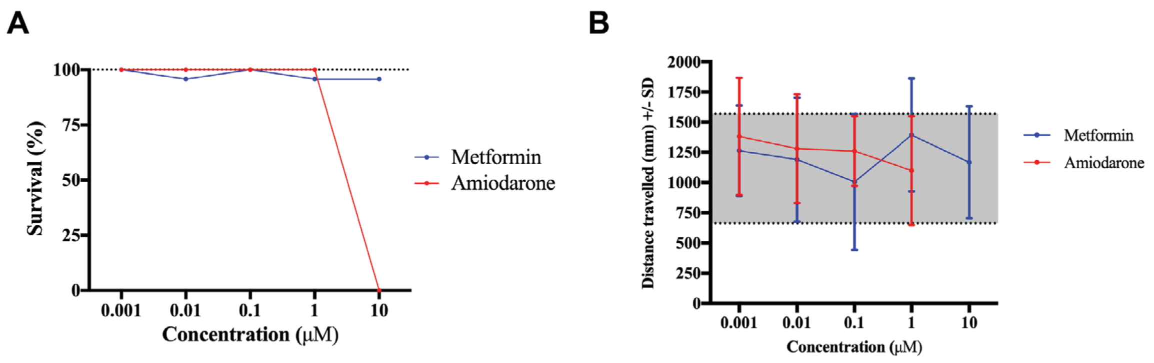


**Supplementary Figure 6:** Evaluation of non-toxic doses of Metformin and Amiodarone for treatment of 6 dpf zebrafish. (A) Effect of different drug concentrations on the survival of fish. (B) Effect of different drug concentrations on swimming capacity.

**Supplementary Table 1:** List of all primers used in the current study

| Primer name | Description | Sequence (5' – 3') |
|---|---|---|
| *bag3* Exon2 gRNA sequence | Gene specific primer for gRNA synthesis. *bag3* (exon 2) targeting sequence is underlined. | TAGG<u>AATGTCCCCAGAGACTCCTC</u> |
| *bag3* exon2 Stop Cassette | Stop cassette primer sequence used for insertion into *bag3* target site | TTCAATGTCCCCAGAGACTCGTCATGGCGTTTAAACCTTAATTAAGCTGTTGTAGCTCAGGACATGCACAAAACC |
| *bag3* genotyping F | Genotyping primer targeting *bag3* intron 1 | ACCCAATTGCAATGGAAAAG |
| *bag3* genotyping R | Genotyping primer targeting *bag3* intron 2 | CCTTTCCCAATAAACCCAAGA |

**Supplementary Table 2:** Number of fish used for experiments presented in Figure 2

| | Biological replicate 1 | Biological replicate 2 | Biological replicate 3 | Total |
|---|---|---|---|---|
| *bag3* $^{+/+}$ (Total) | 12 | 5 | 11 | **28** |
| *bag3* $^{+/-}$ (Unaffected) | 6 | 11 | 10 | **27** |
| *bag3* $^{+/-}$ (Affected) | 3 | 4 | 1 | **8** |
| *bag3* $^{-/-}$ (Unaffected) | 3 | 5 | 3 | **11** |
| *bag3* $^{-/-}$ (Affected) | 4 | 5 | 4 | **13** |

**Supplementary Table 3:** Number of fish, and statistical tests, including t/F values and p values for experiments in presented in Figure 3.

| | | | | | | | | |
|---|---|---|---|---|---|---|---|---|
| **3A: Maximum Acceleration** | | | | | | | | |
| Sibling Control for Tg(BAG3$^{wt}$-eGFP) | 14 | 17 | 19 | | 50 | | t = 1.154; | - |
| Tg(BAG3$^{wt}$-eGFP) | 13 | 18 | 19 | | 50 | t-test | df = 81.464 | 0.251 |
| **3B: Maximum Acceleration** | | | | | | | | |
| Sibling Control for Tg(BAG3$^{P209L}$-eGFP) | 15 | 19 | 17 | | 51 | | t = -1.924 | - |
| Tg(BAG3P209L-eGFP) | 15 | 20 | 18 | | 53 | t-test | df = 92.573 | 0.057 |
| **3C: Distance travelled** | | | | | | | | |
| Sibling Control for Tg(BAG3$^{wt}$-eGFP) | 24 | 24 | 24 | 16 | 88 | | t = -1.123 | - |
| Tg(BAG3$^{wt}$-eGFP) | 24 | 22 | 22 | 22 | 90 | t-test | df = 176 | 0.263 |
| **3D: Distance travelled** | | | | | | | | |
| Sibling Control for Tg(BAG3$^{P209L}$-eGFP) | 22 | 24 | 24 | 22 | 92 | | t = 0.224 | - |
| Tg(BAG3$^{P209L}$-eGFP) | 23 | 23 | 23 | 16 | 85 | t-test | df = 175 | 0.823 |
| **3E: Maximum Acceleration** | | | | | | | | |
| *bag3* $^{+/+}$ | 21 | 14 | 18 | | 53 | One way ANOVA | | - |
| *bag3* $^{+/-}$ | 31 | 33 | 30 | | 94 | with Dunnett's post | F = 0.786; | 0.174 |
| *bag3* $^{-/-}$ | 16 | 13 | 16 | | 45 | hoc test | df = 2 | 0.359 |
| **3F: Distance travelled** | | | | | | | | |
| *bag3* $^{+/+}$ | 23 | 21 | 17 | | 61 | One way ANOVA | | - |
| *bag3* $^{+/-}$ | 47 | 47 | 49 | | 143 | with Dunnett's post | F = 6.924; | 0.000239 |
| *bag3* $^{-/-}$ | 19 | 22 | 15 | | 56 | hoc test | df = 2 | 0.035 |
| **3G: Distance travelled** | | | | | | | | |
| Tg(BAG3$^{wt}$-eGFP), *bag3* $^{+/+}$ | 26 | 11 | 22 | | 59 | One way ANOVA | | - |
| Tg(BAG3$^{wt}$-eGFP), *bag3* $^{+/-}$ | 29 | 27 | 39 | | 95 | with Dunnett's post | F = 0.36; | 0.416 |
| Tg(BAG3$^{wt}$-eGFP), *bag3* $^{-/-}$ | 10 | 6 | 18 | | 34 | hoc test | df = 2 | 0.327 |
| **3H: Distance travelled** | | | | | | | | |
| Tg(BAG3$^{P209L}$-eGFP), *bag3* $^{+/+}$ | 15 | 22 | 20 | | 57 | One way ANOVA | | - |
| Tg(BAG3$^{P209L}$-eGFP), *bag3* $^{+/-}$ | 47 | 49 | 28 | | 124 | with Dunnett's post | F = 0.217; | 0.377 |
| Tg(BAG3$^{P209L}$-eGFP), *bag3* $^{-/-}$ | 24 | 20 | 12 | | 56 | hoc test | df = 2 | 0.516 |

**Supplementary Table 4:** Number of fish used for experiments presented in Figure 6. A one way ANOVA (two tailed) was used to compare differences in aggregate numbers and the resulting p values are noted.

| | Drug | Number of fish/replicate | | | | Mean difference in log(aggregate count) (Drug-DMSO) | Standard error | p value |
|---|---|---|---|---|---|---|---|---|
| | | 1 | 2 | 3 | Total | | | |
| 1 | Rapamycin | 5 | 4 | 5 | **14** | -1.8506 | 0.4180 | 0.000252 |
| 2 | Timosaponin A-III | 0 | 0 | 0 | **0** | | | |
| 3 | PI-103 | 4 | 4 | 5 | **13** | -1.8325 | 0.48961 | 0.004 |
| 4 | Lithium Chloride | 5 | 5 | 4 | **14** | 0.0144 | 0.4180 | 1 |
| 5 | L-690,330 | 5 | 5 | 5 | **15** | -0.0466 | 0.37776 | 1 |
| 6 | Sodium Valproate | 4 | 5 | 5 | **14** | -0.3537 | 0.48046 | 0.999 |
| 7 | Verapamil·HCl | 5 | 4 | 5 | **14** | 0.51460 | 0.39660 | 0.862 |
| 8 | Loperamide·HCl | 3 | 4 | 5 | **12** | -0.4396 | 0.4344 | 0.978 |
| 9 | Amiodarone·HCl | 5 | 5 | 5 | **15** | -2.3736 | 0.47238 | 0.000019 |
| 10 | Nimodipine | 5 | 5 | 5 | **15** | -0.9168 | 0.37776 | 0.151 |
| 11 | Nitrendipine | 5 | 4 | 5 | **14** | -0.1825 | 0.4180 | 1 |
| 12 | Niguldipine | 5 | 4 | 4 | **13** | -1.4956 | 0.48961 | 0.033 |
| 13 | Penitrem A | 5 | 4 | 4 | **13** | 0.51050 | 0.40416 | 0.882 |
| 14 | Ionomycin | 0 | 0 | 0 | **0** | | | |
| 15 | Rotenone | 0 | 0 | 0 | **0** | | | |
| 16 | TTFA | 4 | 5 | 5 | **14** | -0.7138 | 0.4256 | 0.592 |
| 17 | Fluspirilene | 3 | 5 | 5 | **13** | -1.0813 | 0.48961 | 0.258 |
| 18 | Trifluoperazine·2HCl | 4 | 5 | 5 | **14** | 1.34990 | 0.39660 | 0.01 |
| 19 | Sorafenib tosylate | 5 | 5 | 4 | **14** | -0.6559 | 0.4180 | 0.676 |
| 20 | Niclosamide | 5 | 5 | 4 | **14** | 0.1040 | 0.4180 | 1 |
| 21 | Rottlerin | 5 | 5 | 5 | **15** | -0.09130 | 0.38993 | 1 |
| 22 | Caffeine | 5 | 4 | 4 | **13** | 0.095 | 0.48961 | 1 |
| 23 | Metformin·HCl | 4 | 5 | 5 | **14** | -1.6676 | 0.38422 | 0.000344 |
| 24 | Clonidine·HCl | 5 | 4 | 5 | **14** | 0.67770 | 0.39660 | 0.56 |
| 25 | Rilmenidine | 5 | 5 | 5 | **15** | 0.26680 | 0.38993 | 1 |
| 26 | 2',5'-Dideoxyadenosine | 4 | 5 | 5 | **14** | -0.1705 | 0.4180 | 1 |
| 27 | Suramin·6Na | 3 | 4 | 3 | **10** | 0.1886 | 0.4446 | 1 |
| 28 | Pimozide | 4 | 5 | 5 | **14** | 1.28030 | 0.39660 | 0.018 |
| 29 | STF-62247 | 4 | 5 | 5 | **14** | -1.6811 | 0.48046 | 0.008 |
| 30 | Spermidine | 5 | 5 | 5 | **15** | -1.0031 | 0.4112 | 0.151 |
| 31 | FK-866 | 4 | 4 | 5 | **13** | 1.20470 | 0.40416 | 0.038 |
| 32 | Tamoxifen citrate | 5 | 4 | 4 | **13** | -0.5464 | 0.48961 | 0.961 |
| 33 | Minoxidil | 5 | 5 | 5 | **15** | -0.35 | 0.37776 | 0.988 |
| 34 | Imiquimod | 4 | 4 | 4 | **12** | -0.1639 | 0.4344 | 1 |
| 35 | Imatinib mesylate | 3 | 5 | 4 | **12** | 0.53830 | 0.41280 | 0.858 |
| 36 | AG112 | 5 | 5 | 5 | **15** | -1.3655 | 0.37776 | 0.005 |
| 37 | SU11652 | 5 | 5 | 5 | **15** | 1.7976 | 0.47238 | 0.003 |
| 38 | SB202190 | 5 | 5 | 5 | **15** | 0.50260 | 0.38993 | 0.867 |
| 39 | Brefeldin A | 4 | 5 | 5 | **14** | -1.6205 | 0.4180 | 0.002 |
| 40 | Tunicamycin | 0 | 0 | 0 | **0** | | | |
| 41 | Thapsigargin | 0 | 0 | 0 | **0** | | | |
| 42 | A23187 | 0 | 0 | 0 | **0** | | | |
| 43 | Capsaicin | 4 | 5 | 4 | **13** | -0.4420 | 0.4256 | 0.972 |
| 44 | Dihydrocapsaicin | 5 | 5 | 5 | **15** | -0.6361 | 0.37776 | 0.579 |

| | Drug | Number of fish/replicate | | | | Mean difference in log(aggregate count) (Drug-DMSO) | Standard error | p value |
|---|---|---|---|---|---|---|---|---|
| | | 1 | 2 | 3 | Total | | | |
| 45 | Glucosamine HCl | 4 | 5 | 5 | **14** | -0.9986 | 0.38422 | 0.102 |
| 46 | DTT | 4 | 4 | 5 | **13** | 0.1904 | 0.39154 | 1 |
| 47 | Deoxycholate·Na | 4 | 5 | 5 | **14** | -0.8018 | 0.38422 | 0.303 |
| 48 | ABC294640·HCl | 5 | 4 | 5 | **14** | -0.6031 | 0.38422 | 0.668 |
| 49 | Licochalcone A | 3 | 4 | 4 | **11** | -3.7706 | 0.40958 | <0.0001 |
| 50 | Curcumin | 4 | 5 | 5 | **14** | 3.46120 | 0.39660 | <0.0001 |
| 51 | Plumbagin | 0 | 0 | 0 | **0** | | | |
| 52 | 6-Gingerol | 5 | 5 | 3 | **13** | -0.7948 | 0.39154 | 0.336 |
| 53 | Akt Inhibitor X·HCl | 5 | 5 | 4 | **14** | 0.0282 | 0.48046 | 1 |
| 54 | PMSF | 5 | 5 | 4 | **14** | -0.9846 | 0.48046 | 0.348 |
| 55 | MG132 | 4 | 4 | 5 | **13** | 1.18670 | 0.40416 | 0.043 |
| 56 | ALLN | 4 | 5 | 5 | **14** | 0.12470 | 0.39660 | 1 |
| 57 | 7-Ketocholesterol | 5 | 5 | 5 | **15** | -0.2344 | 0.4112 | 1 |
| 58 | 17-AAG | 4 | 5 | 4 | **13** | 0.3894 | 0.4256 | 0.991 |
| 59 | Geldanamycin | 4 | 5 | 5 | **14** | -0.6168 | 0.48046 | 0.894 |
| 60 | C1 | 5 | 4 | 4 | **13** | 0.0215 | 0.48961 | 1 |
| 61 | Z36 | 5 | 5 | 5 | **15** | 0.8522 | 0.37776 | 0.218 |
| 62 | Rockout | 5 | 4 | 4 | **13** | -0.8741 | 0.48961 | 0.532 |
| 63 | Go6850 | 5 | 4 | 5 | **14** | -0.6866 | 0.38422 | 0.5 |
| 64 | 2-Deoxyglucose | 5 | 5 | 5 | **15** | -1.197 | 0.47238 | 0.129 |
| 65 | Etoposide | 4 | 5 | 5 | **14** | -0.9591 | 0.48046 | 0.381 |
| 66 | SMER28 | 4 | 5 | 4 | **13** | -0.4831 | 0.48961 | 0.987 |
| 67 | Trehalose | 5 | 5 | 5 | **15** | 0.97460 | 0.38993 | 0.129 |
| 68 | C2-dihydroceramide | 3 | 5 | 5 | **13** | -0.4936 | 0.4256 | 0.936 |
| 69 | Temozolomide | 4 | 5 | 4 | **13** | -0.7083 | 0.39154 | 0.39154 |
| 70 | Resveratrol | 5 | 4 | 5 | **14** | 0.71190 | 0.39660 | 0.495 |
| 71 | Staurosporine | 0 | 0 | 0 | **0** | | | |
| Control | DMSO | 18 | 18 | 19 | **55** | | | |
| Control | E3 embryo medium | 18 | 20 | 19 | **57** | | | |

**Supplementary Table 5:** Details of the number of fish examined, statistical tests used and resulting statistical output for experiments presented in Figure 8.

| **Number of fish per biological replicate** | | | | | | | | **Significance test used** | **F value and df** | **p value** |
|---|---|---|---|---|---|---|---|---|---|---|
| | **1** | **2** | **3** | **4** | **5** | **6** | **Total** | | | |
| **Figure 8A-C** | | | | | | | | | | |
| *bag3* $^{-/-}$, Untreated | 7 | 10 | 7 | 7 | | | **31** | Figure 8B: Chi-square test | chi-square = 8.315; df = 2 | 0.0156 |
| *bag3* $^{-/-}$, Metformin | 6 | 4 | 6 | 4 | | | **20** | Figure 8C: One way ANOVA with Dunnett's post hoc test | F = 7.334; df = 2 | p = 0.0061 (Metformin treated); p = 0.0019 (Amiodarone treated) |
| *bag3* $^{-/-}$, Amiodarone | 7 | 3 | 6 | 5 | | | **21** | | | |
| **Figure 8D-G: Distance travelled** | | | | | | | | | | |
| *bag3* $^{+/+}$ | 21 | 24 | 28 | 20 | 19 | 28 | **140** | Figure 8D: Two way ANOVA | F = 3.765; df = 4 | Genotype * Drug interaction: p = 0.005 |
| *bag3* $^{+/-}$ | 47 | 43 | 35 | 39 | 39 | 37 | **240** | Figure 8E: One way ANOVA with Dunnett's post hoc test | F = 2.799; df = 2 | p = 0.043 (bag3 $^{+/-}$) |
| *bag3* $^{-/-}$ | 18 | 14 | 21 | 23 | 16 | 12 | **104** | | | p = 0.031 (bag3 $^{+/-}$) |
| *bag3* $^{+/+}$ + Metformin | 19 | 16 | 21 | 18 | 13 | 19 | **106** | Figure 8F: One way ANOVA with Dunnett's post hoc test | F = 7.857; df = 2 | p = 0.000134 (bag3 $^{-/-}$ Metformin treated) |
| *bag3* $^{+/-}$ + Metformin | 44 | 50 | 42 | 31 | 30 | 38 | **235** | | | p = 0.197 (bag3 $^{-/-}$ Amiodarone treated) |
| *bag3* $^{-/-}$ + Metformin | 21 | 17 | 13 | 14 | 24 | 15 | **104** | Figure 8G: One way ANOVA with Dunnett's post hoc test | F = 0.377; df = 2 | p = 0.619 (bag3 $^{-/-}$ Metformin treated) |
| *bag3* $^{+/+}$ + Amiodarone | 18 | 13 | 16 | 15 | 24 | 16 | **102** | | | p = 0.897 (bag3 $^{-/-}$ Amiodarone treated) |
| *bag3* $^{+/-}$ + Amiodarone | 43 | 53 | 50 | 39 | 30 | 43 | **258** | Figure 8H One way ANOVA with Dunnett's post hoc test | F = 7.857; df = 2 | p = 0.059 (bag3 $^{+/+}$ Metformin treated) |
| *bag3* $^{-/-}$ + Amiodarone | 24 | 23 | 14 | 9 | 21 | 15 | **106** | | | p = 0.772 (bag3 $^{+/+}$ Amiodarone treated) |

**Supplementary Table 6:** Number of fish used for experiments presented in Supplementary Figure 6.

| **Supplementary Figure 7: Distance travelled** | |
|---|---|
| | **Total** |
| Untreated | **23** |
| 0.001 µM Metformin | **24** |
| 0.01 µM Metformin | **23** |
| 0.1 µM Metformin | **24** |
| 1 µM Metformin | **23** |
| 10 µM Metformin | **24** |
| 0.001 µM Amiodarone | **24** |
| 0.01 µM Amiodarone | **24** |
| 0.1 µM Amiodarone | **24** |
| 1 µM Amiodarone | **24** |

**Supplementary Table 7:** Patient details for biopsy samples.

| **Experiment** | **Sample** | **Age of onset** | **Age at biopsy** | **Biopsy location** | **Previous publication of patient** |
|---|---|---|---|---|---|
| p62 immunohistochemistry | Control | - | 30 | Deltoid | - |
| | BAG3$^{P209L}$ patient | 11 | 31 | Deltoid | 48 |
| | ZASP$^{A165V}$ distal myopathy | 45 | 54 | Quadriceps | |
| | Myotilin$^{S60F}$ distal myopathy patient | 65 | 71 | *Tibialis anterior* | |
| | Desmin | 63 | 70 | *Peroneus longus* | |
| p62 and LC3 western | Control muscle | - | 18 years | *Biceps Brachialis* | |
| | Control brain | | 48 years | Parietal cortex | |
| | patient muscle | 14 years | 16 years | *Biceps Brachialis* | 49 |
